# Therapeutic Potential of Melatonin, Erythropoietin, and TRPV4 Antagonism in a Preclinical Model of Genetic Hydrocephalus

**DOI:** 10.64898/2026.11.05.764645

**Authors:** Fatemeh Omidi Bidgoli, David Audu, Gowthami Mahendran, Yanping Zhang, Lauren L. Jantzie, Shenandoah Robinson, Bonnie L. Blazer-Yost

**Affiliations:** Hydrocephalus Research Center, Department of Biology, Indiana University, Indianapolis, IN, USA; Department of Pediatrics, Division of Neonatal-Perinatal Medicine, Johns Hopkins University School of Medicine, Baltimore, MD, USA; Department of Neurosurgery, Division of Pediatric Neurosurgery, Johns Hopkins University School of Medicine, Baltimore, MD, USA

**Keywords:** Neuroinflammation, TLR4, NLRP3, periventricular structural alterations, ventriculomegaly

## Abstract

Symptomatic hydrocephalus is a neuropathological disorder of cerebrospinal fluid (CSF) homeostasis characterized by abnormal accumulation of CSF and progressive ventricular enlargement. Current treatment relies primarily on surgical CSF diversion, which is invasive and associated with substantial rates of treatment failure and surgical revisions. Congenital hydrocephalus can be caused by genetic pathologies; for example, a point mutation in the Meckel-Gruber type 3, *Tmem*67, gene produces a congenital hydrocephalus phenotype in rats, characterized by progressive ventriculomegaly. Here, we evaluated a combinatorial cocktail therapeutic strategy consisting of erythropoietin (EPO), melatonin (MLT), and the transient receptor potential vanilloid 4 (TRPV4) antagonist, RN1734 (RN), in the *Tmem*67□□□ rat model of congenital hydrocephalus. Magnetic Resonance Imaging (MRI) demonstrated marked progression of ventriculomegaly in untreated *Tmem*67□□□ rats between postnatal (P) 7 and P15. RN1734 and EPO+MLT+RN1734 (triple therapy) from P7-P14 prevented further ventricular expansion. Hydrocephalus is associated with structural disruption of the periventricular white matter (WM) and ependymal lining (EL), accompanied by regionally increased TLR4 and NLRP3-associated inflammatory signaling. EPO+MLT, RN1734, or triple therapy differentially attenuated these inflammatory responses. The triple combination reduced both TLR4 and NLRP3 protein abundance within the affected periventricular region and ameliorated the structural alterations. These findings support a combined therapeutic approach that addresses ventricular pathology, inflammation and periventricular structural consequences of congenital hydrocephalus.

## Introduction

Symptomatic hydrocephalus is a neurological condition resulting from disrupted cerebrospinal fluid (CSF) microcirculatory homeostasis leading to abnormal accumulation of CSF and progressive enlargement of the cerebral ventricular system [17]. Hydrocephalus affects individuals across the lifespan [6, 8]. Depending on its origin, hydrocephalus can be broadly categorized as congenital or acquired [38]. The incidence of congenital hydrocephalus varies by global region, ranging from 316 per 100,000 births in Latin America to 145 per 100,000 births in Africa to 68 per 100,000 births in the United States and Canada [8]. Genetic factors are estimated to contribute to up to 40% of congenital hydrocephalus cases worldwide, though the precise underlying etiology remains unidentified in the majority of individual cases [28]. A recognized subset of genetically driven congenital hydrocephalus arises from mutations in genes that govern ciliary structure and function. Among these is *TMEM67*□□□, which encodes transmembrane protein 67 (meckelin). *TMEM67*□□□ is the most frequently mutated gene in Meckel–Gruber syndrome (MKS), accounting for more than 15% of MKS diagnoses across broad patient populations [1].

Existing treatment strategies for hydrocephalus include several surgical procedures; placement of a ventriculoperitoneal (VP) shunt, endoscopic third ventriculostomy (ETV), and choroid plexus cauterization (CPC) [29]. Despite advances in surgical techniques and shunt designs, shunt failure is common in pediatric and adult patients [10, 21]. These limitations in surgical treatment have highlighted the urgent need for new, non-invasive therapeutic strategies, reflecting a shared priority among patients and the hydrocephalus research community to reduce complications and improve long-term outcomes [2, 7, 16, 33].

Preclinical studies have begun to explore pharmacological alternatives for hydrocephalus treatment. In a rat model of posthemorrhagic hydrocephalus of prematurity (PHHP), combined treatment with erythropoietin (EPO) and melatonin (MLT) reduced macrocephaly and ventriculomegaly, preserved ependymal cilia, and improved neurobehavioral outcomes [39]. Separately, in the *Tmem*67□□□ genetic rat model of congenital hydrocephalus, which is orthologous to Meckel-Gruber syndrome, type 3 [13, 43], antagonism of the mechanosensitive ion channel called Transient Receptor Potential Vanilloid 4 (TRPV4) reduced ventriculomegaly [18] and attenuated Toll-like receptor 4 (TLR4) and NOD-like receptor family pyrin domain-containing 3 (NLRP3)-mediated neuroinflammation and oxidative stress in cingulate cortex [3]. The ventriculomegaly-reducing effect of RN1734 was further supported by complementary *in vitro* studies in human choroid plexus epithelial cells, suggesting that TRPV4 plays an important role in CSF production [19]. EPO is neuroprotective through various mechanisms. Acutely, it exerts anti-apoptotic, anti-inflammatory, neurotrophic, and anti-oxidant effects [27, 36]. Over an extended period of use, EPO promotes brain repair in neural cells [24, 25, 42, 46, 48], including neurons [9, 26, 31], astrocytes [15, 44], and oligodendrocytes [5, 23, 25]. Similarly, MLT is a neuroregulatory hormone with strong antioxidant, free radical scavenging, anti-inflammatory, and immunomodulatory properties, which may contribute to its effects on neurological disease pathophysiology [14, 20]. It also boosts immune responses to pathogens [4] and limits inflammatory tissue damage through direct free radical scavenging and indirect suppression of damage-associated mediators [11, 37]. MLT has additionally shown therapeutic promise across a range of conditions, including depression, aging, rheumatoid arthritis, fibromyalgia, chronic fatigue syndrome, gastrointestinal and blood disorders, ocular disease, and cardiovascular disease [40].

Hydrocephalus is a multifactorial disorder involving abnormal CSF accumulation and ventricular enlargement together with secondary structural injury, neuroinflammation, oxidative stress, and cellular damage [3, 17, 47, 49]. A therapeutic approach directed at a single pathological process may therefore provide incomplete protection. In the present study, we use the *Tmem*67□□□ rat model of genetic hydrocephalus to compare the RN1734 (TRPV4 antagonist), EPO and MLT (EPO+MLT), or triple combination therapy (EPO+MLT+RN). The rationale for the triple combination therapy was to target multiple pathological consequences of hydrocephalus through complementary therapeutic actions. This approach compares the potential benefits of triple combination therapy relative to individual or dual treatment approaches.

## Materials and methods

### Animals and Genotyping

In this study, we used outbred Wistar wild-type rats in addition to our inbred wild-type (*Tmem*67□□□) and homozygous (*Tmem*67□□□) animals. Outbred Wistar rats were obtained from Dr. Nicholas Grahame (Department of Psychology, Indiana University Indianapolis, Indiana, USA) and used as an additional reference control. The comparison between Wistar and *Tmem*67□□□ animals was used to determine whether the inbred line demonstrated differences in the parameters measured in this study, while the comparison between *Tmem*67□□□ and *Tmem*67□□□ animals shows the effect of treatment in the *Tmem*67 line.

For the inbred colony, heterozygous *Tmem*67□□□ rats were bred to produce litters containing *Tmem*67□□□, *Tmem*67□□□, and *Tmem*67□□□ offspring. The rat colony was originally provided by Dr. Vincent Gattone (Department of Anatomy and Cell Biology, Indiana University School of Medicine). For experiments, *Tmem*67□□□ pups and age-matched *Tmem*67□□□ littermates were treated daily starting at postnatal (P) 7 and continuing to P14. Tissue was collected at P15 (Fig. 1a). Both male and female animals were included in the study. Animals were selected for experiments in a blinded manner, and animals that died before reaching P15 (2 animals treated with RN) were also included in the study.

**Fig. 1.**
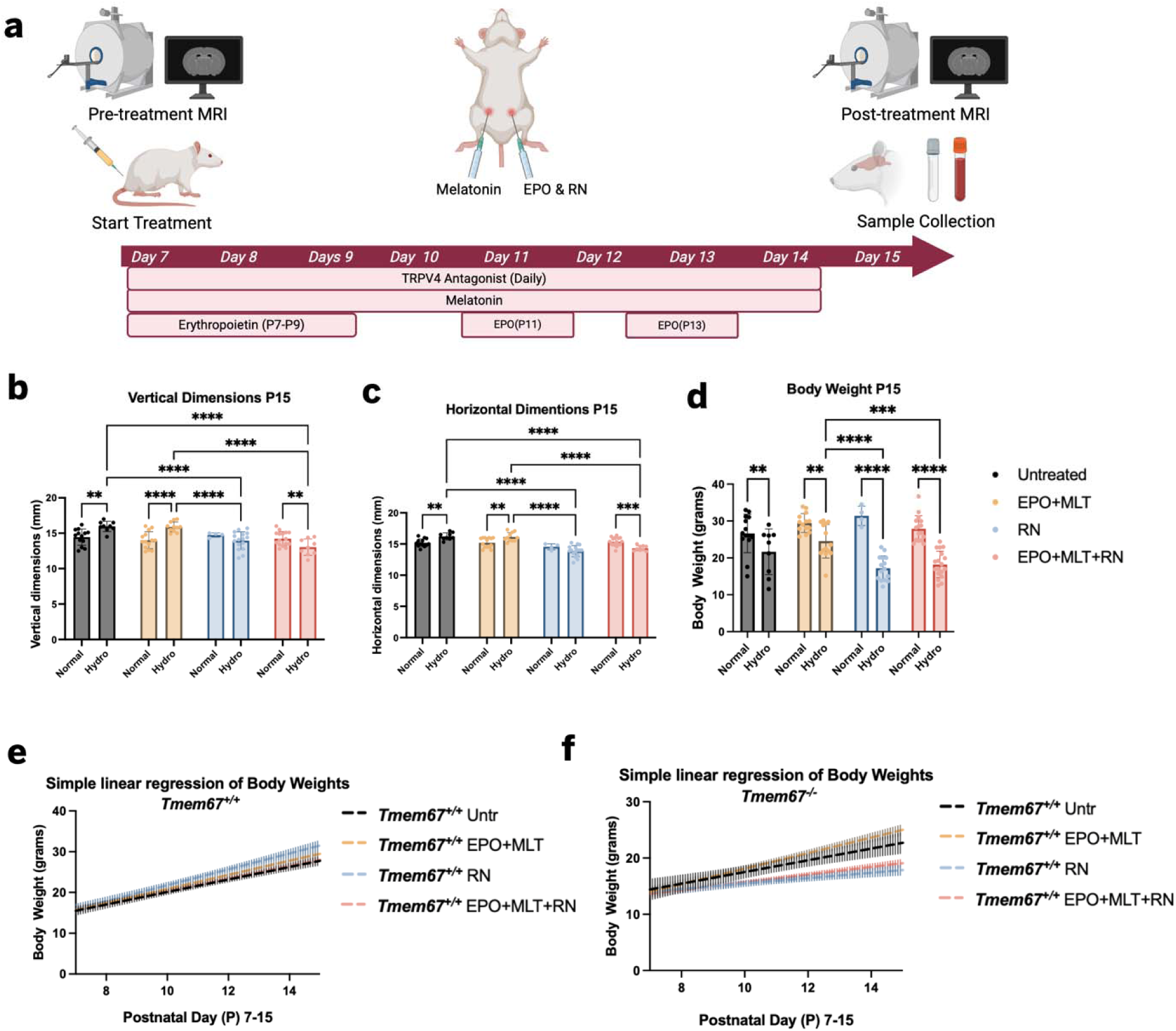
Experimental design and effects of triple combination therapy on cranial dimensions and body weight in a genetic rat model of hydrocephalus. (a) Schematic overview of the triple combination therapy protocol in a congenital model of hydrocephalus. Pre-treatment MRI was performed prior to treatment initiation on P7. Animals received daily i.p. injections of the TRPV4 antagonist, RN1734 (4 mg/kg B.W.) and MLT (20 mg/kg B.W.) from P7 through P14, along with EPO (1000 U/kg) administered on P7–P9, P11, and P13. RN1734 and EPO were injected on the left side of the animals, and MLT was injected on the right side. Post-treatment MRI and sample collection (blood, CSF, and tissue (CPX, CTX, WM, SVZ)) were performed on P15. (b, c) Vertical (b) and horizontal (c) cranial dimensions of wildtype (*Tmem*67□□□) and homozygous (*Tmem*67□□□) animals, untreated or treated with EPO+MLT, RN1734, or triple combination therapy (EPO+MLT+RN1734), measured at P15, one day after the final treatment. Statistical significance was determined by two-way ANOVA with Tukey’s multiple comparisons test. Data in (b, c) are presented as mean ± SEM (Untreated: n=13 Normal, n=9 Hydro; EPO+MLT Treated: n=14 Normal, n=11 Hydro; RN1734 Treated: n=4 Normal, n=16 Hydro; EPO+MLT+RN1734 Treated: n=16 Normal, n=9 Hydro). (d) Body weight of pups at P15, following 8 days of treatment with EPO+MLT, RN1734, or EPO+MLT+RN1734. Data were analyzed by two-way ANOVA with Tukey’s multiple comparisons test. Data in (d) are presented as mean ± SEM (Untreated: n=14 Normal, n=9 Hydro; EPO+MLT Treated: n=14 Normal, n=14 Hydro; RN1734 Treated: n=4 Normal, n=15 Hydro; EPO+MLT+RN1734 Treated: n=16 Normal, n=18 Hydro). *p<0.05, **p<0.01, ***p<0.001, ***p<0.0001; ns, not significant. (e, f) Simple linear regression of body weight over time (P7–P15) for *Tmem*67□□□ (e) and *Tmem*67□□□ (f) animals under each treatment condition. In *Tmem*67□□□ animals, regression slopes differed significantly among treatment groups (F(3,538) = 13.99, p<0.0001). In *Tmem*67□□□ animals, slopes did not differ significantly among treatment groups (F(3,415) = 2.152, p=0.0931); a pooled slope of 1.626 was calculated. Elevations (intercepts) differed significantly among *Tmem*67□□□ treatment groups (F(3,418) = 6.720, p=0.0002).

Genotyping was performed by PCR of tail DNA to distinguish *Tmem*67□□□, *Tmem*67□□□, and *Tmem*67□□□ offspring from the inbred colony, as previously described [13]. Briefly, tail snips were collected from pups at P2, and DNA was extracted using the QIAGEN QIAamp DNA Mini Kit (Cat. #51306). PCR was performed using primers flanking the mutation site in exon 12 of *Tmem*67□□□, and purified PCR products were sequenced to identify the C1186T point mutation distinguishing mutant animals.

### Treatment Conditions

Wildtype (*Tmem*67□□□), mutant (*Tmem*67□□□), and outbred Wistar rats were randomly assigned to one of four treatment groups: untreated, EPO+MLT-treated, RN1734-treated, or EPOMLT/RN1734-treated (triple combination therapy).

Erythropoietin (EPO; Bio-Techne, Cat. #287-TC-500) was reconstituted in sterile saline and administered at 1000 U/kg body weight via intraperitoneal (i.p.) injection on P7, P8, P9, P11, and P13, on the left lower abdominal quadrant of the animal. Injection volume was calculated as body weight (g) × 2 μL/gr of stock solution (500 U/mL).

Melatonin (MLT; Sigma, Cat. #M5250) was prepared in sterile saline at 2 mg/mL and administered at 20 mg/kg body weight via i.p. injection daily from P7 to P14, on the right lower abdominal quadrant of the animal. Injection volume was calculated as body weight (g) × 10 μL/gr of stock solution.

RN1734 (Tocris, Cat. #3746), a TRPV4 antagonist, was prepared in DMSO at 2 mg/mL and administered at 4 mg/kg body weight via i.p. injection daily from P7 to P14, on the left lower abdominal quadrant of the animal. Injection volume was calculated as body weight (g) × 2 μL/gr of stock solution.

### Cranial Measurements

Vertical and horizontal head measurements were taken using calipers before and after drug treatment (P7 and P15) in treated and untreated wildtype (*Tmem*67□□□) and homozygous (*Tmem*67□□□) animals. Vertical measurements were taken from the top of the head to the jawline, and horizontal measurements were taken from ear to ear. All measurements were performed by a single investigator blinded to treatment group to ensure consistency.

### Magnetic Resonance Imaging (MRI)

MRI scans were performed using a 1T horizontal bore M7 MRI system (Aspect Imaging, Shoham, Israel), with a 23 mm coil. Before being placed in the coil, rats were anesthetized using isoflurane (Pivetal; induction 4%, maintenance 2%) delivered via an R550 tabletop veterinary anesthesia machine (RWD Life Science, USA). Core temperature was maintained at approximately 37°C using warm water flow under the bed of the coil, and respiratory rate was monitored using PC-SAM monitoring software throughout the scan.

Anatomical T2-weighted images were acquired through the brain using a 3D fast spin echo (FSE) sequence with the following parameters: repetition time (TR) 3000 ms, echo time (TE) 176.85 ms (≈180 ms), inversion time (TI) 100 ms, echo train length (ETL) 32, echo spacing 9.31 ms, 2 excitations/averages, 64 phase encodings, field of view 20 mm (horizontal) × 30 mm (vertical), 30 slices, slice thickness 0.4 mm (400 µm), 0 mm inter-slice gap, and in-plane resolution 0.31 × 0.3 mm (≈300 µm), yielding a final voxel resolution of 300 × 300 × 400 µm³. Average total scan time for T2-weighted imaging was approximately 8 minutes, during which respiratory rate was continuously monitored.

After scan completion, rat pups were removed from the coil and allowed to recover for about 3–5 minutes, after which animals were returned to their home cages. Scans were quantified using Imalytics Preclinical software (Version 3.1.3; Gremse-IT GmbH, Germany). All scans were analyzed by an investigator blinded to treatment conditions and genotypes.

### Tissue preparation

At P15, animals were deeply anesthetized with isoflurane and transcardially perfused with 0.1 M phosphate-buffered saline (PBS), followed by 4% paraformaldehyde (PFA) prepared in 0.1 M PBS. Brains were collected, post-fixed overnight in 4% PFA at 4°C and cryoprotected in 30% sucrose until fully submerged. Tissues were then embedded in OCT compound, flash-frozen, and coronally sectioned at 15 µm using a cryostat. The region of interest was defined as the pericallosal corpus callosum, using the *Atlas of the Neonatal Rat Brain* [35] as an anatomical reference. Sections corresponding approximately to the P14 coronal plate region C10–C12, in which the lateral ventricle and pericallosal corpus callosum were clearly identifiable, were selected for analysis. Three representative adjacent coronal sections per animal containing the defined region of interest were selected for immunofluorescence and Hematoxylin and Eosin (H&E) staining.

### Immunofluorescence (IF) Staining

Slide-mounted coronal brain sections were washed in PBS. Antigen retrieval was performed in 10 mM sodium citrate buffer (pH 6.0; 2.94 g trisodium citrate dihydrate in 1 L DDI H O, with 0.5 mL Tween-20 added). Sections were brought to a boil and maintained at a sub-boiling temperature for 10 min, followed by cooling at room temperature for 30 min while submerged in the retrieval buffer. After washing in PBS, sections were permeabilized with 1% Triton X-100 before blocking. Sections were subsequently blocked for 1 h at room temperature in blocking buffer containing 1% bovine serum albumin (BSA) in PBS and incubated overnight at 4°C with the appropriate primary antibodies (Table 1). Following PBS washes, sections were incubated for 1 h at room temperature with the appropriate species-specific secondary antibodies (Table 1). Nuclei were counterstained with 4′,6-diamidino-2-phenylindole dihydrochloride (DAPI; Sigma-Aldrich, Cat. #D9542). Following final washes, sections were coverslipped using Aqua-Poly/Mount (Polysciences, Inc.) and protected from light until imaging.

**Table 1.** Antibodies Used in This Study.

| Target / Antibody | Host Species | Dilution | Manufacturer | Catalog # | Application |
|---|---|---|---|---|---|
| **Primary Antibodies (IF)** |  |  |  |  |  |
| TLR4 | Rabbit | 1:100 | Novus Biologicals | NB100-56580 | IF |
| NLRP3 | Rabbit | 1:100 | Novus Biologicals | NBP2-12446SS | IF |
| **Secondary Antibodies (IF)** |  |  |  |  |  |
| Anti-Rabbit IgG (AF546) | Goat anti-rabbit | 1:1,000 | Invitrogen | A-11035 | IF |
| **Primary Antibodies (WB)** |  |  |  |  |  |
| NLRP3 | Rabbit | 1:500 | Novus Biologicals/Bio-Techne | NBP2-12446 | WB |
| TLR4 | Rabbit | 1:300 | Novus Biologicals/Bio-Techne | NB100-56580 | WB |
| β-Actin (loading control) | Mouse | 1:5,000 | Proteintech | 66009-1-Ig | WB |
| **Secondary Antibodies (WB)** |  |  |  |  |  |
| HRP Anti-Rabbit | Anti-Rabbit | 1:10,000 | Invitrogen | 31460 | WB |
| HRP Anti-Mouse | Anti-Mouse | 1:10,000 | Invitrogen | 31430 | WB |
| StrepTactin-HRP (MW standard detection) | StrepTactin-HRP Conjugate | 1:3,000 | Bio-Rad | 1610381 | WB |

### Hematoxylin and Eosin (H&E)

General brain tissue morphology was assessed using an H&E staining kit (Novus Biologicals, Cat. #H-3502-NB). Brain sections (15 µm) were rinsed in distilled deionized water and then stained with Mayer’s hematoxylin for 5 min, rinsed in distilled water, and treated with bluing reagent for 15 s. Following two additional washes, sections were briefly immersed in 100% ethanol for 10 sec and counterstained with Eosin Y for 3 min. Sections were subsequently rinsed and dehydrated through three changes of 100% ethanol, cleared in xylene, and coverslipped using a permanent mounting medium. Stained sections were examined for histological alterations in brain tissue architecture, with particular attention to the ventricular and periventricular regions.

### Image Acquisition and Analysis

Images were acquired using microscopy systems appropriate for each staining modality. H&E sections were imaged using a KEYENCE BZ-X800 All-in-One Fluorescence Microscope under brightfield illumination. Immunofluorescence (IF) was assessed using a Nikon AX R laser-scanning confocal microscope. Acquisition settings, including laser power, gain, and pinhole size, were maintained consistently across experimental groups to permit quantitative comparison of fluorescence signals. Image-based protein-associated fluorescence intensity was quantified using ImageJ/Fiji software (NIH, version 1.54p). Mean gray value, defined as the average pixel intensity within a specified region of interest (ROI) was measured for each image. Background fluorescence was corrected using the rolling-ball background subtraction method to minimize non-specific signals. Fluorescence intensity values are reported in arbitrary units (A.U.) and were used for relative comparisons of protein immunoreactivity across experimental groups.

### Western Blotting

Tissue samples were homogenized in Pierce™ RIPA Buffer (Cat. #89901; Thermo Scientific, Rockford, IL, USA) using a sonicator (Fisher Scientific, Pittsburgh, PA, USA) with two 10-second pulses at 20% and 50% power, respectively. Samples were returned to ice for several minutes between sonication pulses to minimize heating. The resulting homogenates were incubated on ice for 30 min and subsequently centrifuged at 21,100 × *g* for 20 min at 4 °C. The supernatant was carefully transferred to a new tube, and the pellet was discarded. Samples were maintained on ice for immediate analysis or stored at −80 °C until protein concentration determination.

Protein concentration was determined using a detergent-compatible (DC) protein assay (Bio-Rad, Hercules, CA) according to the manufacturer’s protocol. The samples were denatured at 95 °C for 5 minutes and separated on gradient (4–15%) Stain-Free precast midi gels (Cat. #5678084, Bio-Rad) at 150V. Proteins were transferred using a Trans-Blot Turbo system (Bio-Rad) at 25V, 2.5A for 7 minutes using the mixed molecular weight protocol.

Membranes were incubated overnight at 4 °C with primary antibody against NLRP3 (rabbit, 1:500; Novus Biologicals/Bio-Techne, Cat. #NBP2-12446; ∼110–120 kDa), followed by an incubation with HRP-conjugated goat-anti-rabbit secondary antibody (1:10,000; Invitrogen, Cat. #31460) for 1 hour at room temperature. The blotting signals were visualized by ECL substrate (Bio-Rad,Cat. #102032464) in 1:1 ratio. The membrane was incubated in substrate solution for 5 min prior to imaging with a ChemiDoc™ Imaging System using Image Lab™ Touch Software (version 2.3.0.07; Bio-Rad Laboratories, Hercules, CA, USA). Membranes were subsequently stripped and re-probed for TLR4 (rabbit, 1:300; Novus Biologicals/Bio-Techne, Cat. #NB100-56580; ∼95.7 kDa) overnight at 4°C, followed by the HRP-conjugated secondary antibody, ECL substrate, and imaging steps described above. For molecular weight standards, Precision Plus Protein HRP Standards (Bio-Rad, CA. Cat. #1610376) were detected using StrepTactin-HRP conjugate (Cat. #1610381; 1:3000 dilution). The protein expressions were normalized by β-actin (mouse,1:5000, Proteintech) (Table 1).

### Cell culture

HIBCPP cells, a well-characterized human choroid plexus epithelial cell line derived from a female cancer patient [22], were cultured and maintained as previously described [19, 30]. Briefly, cells were maintained in Dulbecco’s Modified Eagle Medium (DMEM) complete medium supplemented with 10% fetal bovine serum (Gemini Bio Benchmark, Cat # 100-106), 1% antibiotic (Penicillin/Streptomycin 100×, Cytiva, Cat No: SV30010), 0.125% insulin (Human recombinant 4 mg/ml in zinc solution, Gibco, Cat No: 12585-014), and 3.7 g/L sodium bicarbonate (Fisher Scientific, Cat No: S233-500) at 37° C and 5% CO, with passaging via 0.25% trypsin–EDTA up to seven passages post-thaw from liquid nitrogen. For electrophysiological experiments, cells were seeded onto Transwell inserts and cultured for 10–14 days, transitioning from complete to serum-free media (DMEM powder, 1% antibiotic (Penicillin/Streptomycin 100×), and 0.125% insulin (Human recombinant 4 mg/ml in zinc solution) and 1.85 g/L sodium bicarbonate) after day 8.

### Electrophysiological studies

An eight-chamber electrophysiology instrument (Physiologic Instruments Inc.) was used to measure transepithelial electrical resistance (TEER), which is inversely proportional to conductance and reflects transepithelial permeability, while simultaneously measuring short-circuit current (Isc) as an indicator of net electrogenic ion transport across the epithelium. A well characterized human choroid plexus epithelial cell line, the HIBCPP cells were used in these experiments [19, 22]. Briefly, HIBCPP monolayers grown on Transwell inserts containing permeable filter wells for 10 to 14 days were equilibrated in serum-free DMEM for 1 hour at 37° C and 5% CO before mounting in the Ussing-chamber electrophysiology system. Prior to recording from cell-seeded inserts, the background resistance of the electrophysiology setup was determined by the system using cell-free membrane inserts. Transwell inserts containing confluent epithelial monolayers were then mounted in chamber sliders (surface area of 1.26 cm²). Both chamber compartments (apical and basolateral sides of the membrane) were filled with freshly prepared, prewarmed, oxygenated serum-free DMEM and maintained at 37 °C using a temperature-controlled water bath. The chambers were continuously aerated with a gas mixture containing 95% O and 5% CO. Electrodes were connected to a multichannel voltage/current-clamp controller, and the system was calibrated before data acquisition. Resistance measured across cell-free membranes was used to correct background resistance and system asymmetry. Cell monolayers were allowed to equilibrate in the chambers for 10 min before recordings were initiated.

Isc (µA/cm^2^), TEER, and conductance (mS/cm^2^) were recorded using AQUIRE 3.0 software (Physiologic Instruments Inc.). TEER was calculated as Rt × A, where Rt represents the measured transepithelial resistance and A represents the surface area of the cell monolayer. Conductance is the inverse of TEER. Short-circuit current and conductance were normalized to surface area and expressed as Isc/A and Gt/A, respectively. Rt was determined using brief current pulses (±10 µA), and resistance was expressed as Ω·cm². Only monolayers with TEER values >300 Ω·cm² were included in experiments. To determine whether EPO and MLT modulate TRPV4-agonist (GSK1016790A) stimulated changes in transepithelial electrolyte flux and barrier permeability, monolayers were treated with EPO and MLT bilaterally (apical and basolateral) with 10 U/ml EPO and 50 µM MLT, or vehicle (PBS), under two treatment paradigms. For short-term treatment, EPO+MLT or vehicle were applied for 10 minutes prior to stimulation with the TRPV4 agonist GSK1016790A (15 nM). For long-term treatment, monolayers were pretreated with EPO+MLT or vehicle for 24 hours prior to TRPV4 stimulation at 37° C. In both paradigms, Isc and conductance were recorded continuously for 30 min after TRPV4 agonist addition. Statistical significance was determined using multiple Student’s t-tests in GraphPad Prism 8.0.1.

### Statistical analysis

Data were analyzed with GraphPad Prism, version 11.1.0, except for electrophysiology experiments, which were analyzed using GraphPad Prism, version 8.0.1. Data are presented as mean ± standard error of the mean (SEM). Two-way ANOVA with Tukey’s multiple comparisons test was used for cranial measurement, body weight, and MRI analyses. One-way ANOVA with Tukey’s multiple comparisons test was used for Western blot and immunofluorescence analyses. Multiple Student’s t-tests were used for electrophysiology experiments. Statistical significance is indicated as follows: *p < 0.05, **p < 0.01, and ***p < 0.001, compared to the indicated group.

## Results

### Triple combination therapy ameliorates the increase in cranial dimensions in homozygous (*Tmem*67□□□) rats

Figure 1a depicts the treatment protocol. Cranial enlargement is a characteristic phenotypic feature, common in neonatal, congenital, and infantile models of hydrocephalus and has been used previously to distinguish hydrocephalic *Tmem*67□□□ rats from *Tmem*67□□□ littermates [41]. Vertical (Fig. 1b) and horizontal (Fig. 1c) cranial dimensions were assessed at P7 and P15, corresponding to the start and end points of treatment, respectively (Fig. 1a). At P15, untreated *Tmem*67□□□ rats exhibited significantly greater vertical cranial dimensions (15.96 ± 0.24 mm) compared with untreated *Tmem*67□□□ littermates (14.45 ± 0.32 mm; p = 0.0014; Fig. 1b). Similarly, horizontal cranial dimensions were significantly greater in untreated *Tmem*67□□□ rats (16.22 ± 0.19 mm) than in *Tmem*67□□□ controls (15.19 ± 0.17 mm; p = 0.0010; Fig. 1c). The same pattern was observed in animals treated with EPO+MLT, with *Tmem*67□□□ rats exhibiting significantly greater vertical (15.89 ± 0.21 mm; p < 0.0001) and horizontal (16.12 ± 0.22 mm; p = 0.0016) cranial dimensions compared with EPO+MLT-treated *Tmem*67□□□ rats (14.01 ± 0.32 mm and 15.19 ± 0.21 mm, respectively). In contrast, following RN1734 treatment, cranial dimensions did not differ significantly between *Tmem*67□□□ and *Tmem*67□□□ rats, with vertical measurements of 14.75 ± 0.16 mm and 13.95 ± 0.31 mm, respectively (p = 0.1761), and horizontal measurements of 14.55 ± 0.24 mm and 13.86 ± 0.22 mm, respectively (p = 0.0834). Following triple combination therapy, *Tmem*67□□□ rats exhibited significantly smaller vertical (13.03 ± 0.38 mm; p = 0.0075) and horizontal (14.32 ± 0.14 mm; p = 0.0007) cranial dimensions compared with similarly treated *Tmem*67□□□ rats (14.23 ± 0.25 mm and 15.35 ± 0.17 mm, respectively; Fig. 1b, c).

### Body weight during disease progression and treatment

Body weights were assessed (Fig. 1d) at P15. As expected, the untreated *Tmem*67□□□ animals had significantly lower body weight than *Tmem*67□□□ littermates. None of the three treatments reversed the body weight differences.

Hydrocephalic animals have lower body weights before starting treatment, therefore we did a simple linear regression analysis of body weight from P7 to P15. Treatment did not significantly affect growth rate (slope) in *Tmem*67□□□ animals (F = 2.152, DFn = 3, DFd = 415, p = 0.0931; Fig. 1e). In *Tmem*67□□□ animals, growth rate (slope) differed significantly among treatment groups (F = 13.99, DFn = 3, DFd = 538, p < 0.0001; Fig. 1f), with EPO+MLT-treated animals showing the numerically highest slope (1.378) compared to untreated animals (1.039).

### Triple combination therapy attenuates ventricular expansion in *Tmem*67□□□ rats

Ventriculomegaly is one of the most significant defining symptoms of hydrocephalus. Ventricular volume was assessed by MRI at P7 and P15 in Wistar, *Tmem*67□□□, and *Tmem*67□□□ rats across untreated, EPO+MLT-, RN1734-, and EPO+MLT+RN1734-treated groups (Fig. 2). Representative MRI images demonstrate ventricular morphology across treatment groups and the range of ventricular expansion observed within each genotype (Fig. 2a– e).

**Fig. 2.**
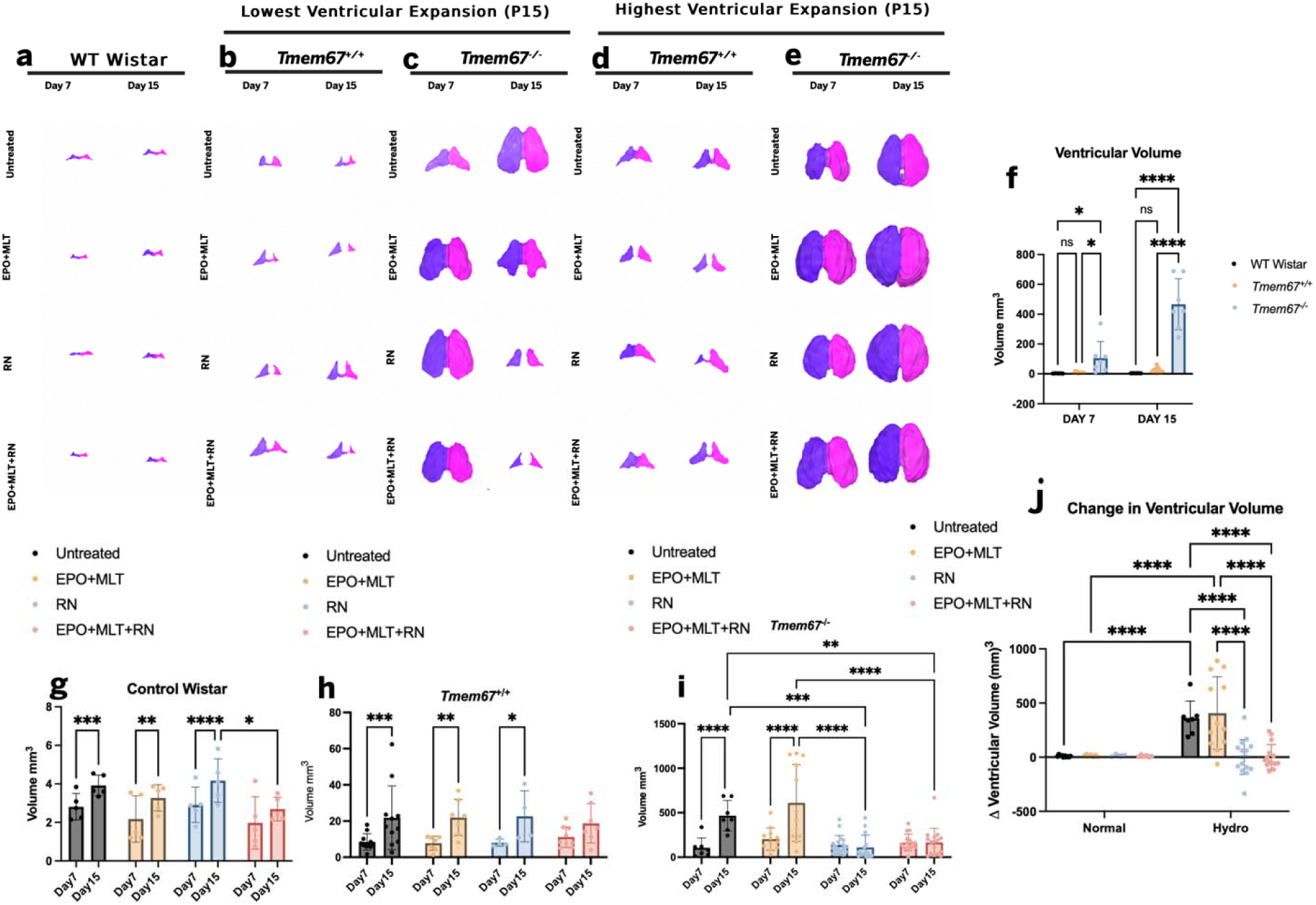
Representative MRI-based 3D reconstructions and quantitative measurements of ventriculomegaly amelioration in hydrocephalic *Tmem*67□□□ rats following treatment. (a–e) Representative 3D reconstructions of brain ventricles (purple, left hemisphere; pink, right hemisphere) at Day 7 and Day 15 in outbred WT Wistar (a), inbred wildtype *Tmem*67□□□ (b, d), and homozygous *Tmem*67□□□ (c, e) rats. Each row corresponds to a different treatment condition (untreated, EPO+MLT, RN1734, or EPO+MLT+RN1734), and each column corresponds to a timepoint (Day 7 or Day 15). For each treatment group, the representative animal with the lowest (b, c) or highest (d, e) ventricular expansion at P15 is shown. (f) Ventricular volume at Day 7 and Day 15, compared across Control Wistar and *Tmem*67□□□ groups. Statistical significance was determined by two-way ANOVA with Tukey’s multiple comparisons test. (g–i) Ventricular volume at Day 7 vs. Day 15, shown separately for Control Wistar (g), *Tmem*67□□□ (h), and *Tmem*67□□□ (i) animals, broken out by treatment group (untreated, EPO+MLT, RN1734, EPO+MLT+RN1734). Statistical significance was determined by two-way ANOVA with Tukey’s multiple comparisons test. (j) Change (Δ) in ventricular volume from Day 7 to Day 15 in Normal vs. Hydrocephalic animals across treatment groups; Normal animals were derived from the *Tmem*67□□□ colony rather than WT Wistar. Statistical significance was determined by two-way ANOVA with Tukey’s multiple comparisons test. Data in (f–j) are presented as mean ± SEM. (f) WT Wistar: Day 7 n=5, Day 15 n=5; *Tmem*67□□□: Day 7 n=12, Day 15 n=12. (g) Control Wistar: untreated n=5 (Day7)/5 (Day15); EPO+MLT n=5/5; RN n=5/5; EPO+MLT+RN1734 n=5/5. (h) *Tmem*67□□□: untreated n=11/11; EPO+MLT n=7/7; RN n=4/4; EPO+MLT+RN1734 n=9/7. (i) *Tmem*67□□□: untreated n=7/7; EPO+MLT n=12/12; RN n=18/15; EPO+MLT+RN1734 n=16/16. (j) Normal: untreated n=10, EPO+MLT n=7, RN n=4, EPO+MLT+RN1734M n=6; Hydro: Non Treated n=7, EPO+MLT n=12, RN n=16, EPO+MLT+RN1734n=15. *p<0.05, **p<0.01, ***p<0.001, ****p<0.0001; ns, not significant.

At P7, ventricular volume did not differ significantly between Wistar and *Tmem*67□□□ rats (2.81 ± 0.31 mm³ and 8.22 ± 1.28 mm³, respectively; *p* = 0.8968). At P15, mean ventricular volume was higher in *Tmem*67□□□ rats (21.59 ± 4.83 mm³) than in Wistar controls (3.92 ± 0.24 mm³), but this difference was not statistically significant (*p* = 0.6716; Fig. 2f). In contrast, *Tmem*67□□□ rats had significantly larger ventricular volume than both Wistar and *Tmem*67□□□ rats at P7 (105.37 ± 41.80 mm³; *p* = 0.0296 and *p* = 0.0120, respectively), and this difference became more pronounced at P15 (466.31 ± 64.97 mm³; *p* < 0.0001 for both; Fig. 2f).

In Wistar rats, ventricular volume increased significantly from P7 to P15 in the untreated (2.81 ± 0.31 to 3.92 ± 0.24 mm³; p = 0.0004), EPO+MLT-treated (2.18 ± 0.54 to 3.27 ± 0.31 mm³; p = 0.0051), and RN1734-treated (2.91 ± 0.41 to 4.17 ± 0.50 mm³; p < 0.0001) groups. No significant increase was observed following EPO+MLT+RN1734 combination treatment (1.98 ± 0.61 to 2.69 ± 0.28 mm³; p = 0.1903; Fig. 2g). A similar pattern was observed in *Tmem*67□□□ rats. Ventricular volume increased significantly between P7 and P15 in untreated (8.50 ± 1.36 to 21.82 ± 5.29 mm³; p = 0.0009), EPO+MLT-treated (7.75 ± 1.47 to 21.97 ± 3.78 mm³; p = 0.0037), and RN1734-treated (8.00 ± 0.97 to 22.66 ± 7.04 mm³; p = 0.0195) animals, whereas the increase following triple combination therapy did not reach statistical significance (11.18 ± 1.91 to 18.72 ± 4.09 mm³; p = 0.0779; Fig. 2h).

In *Tmem*67□□□ animals, ventricular volume increased significantly from P7 to P15 in untreated animals (105.4 ± 41.8 to 466.3 ± 65.0 mm³; p < 0.0001) indicating severe disease progression. This was not diminished in EPO+MLT-treated animals (203.0 ± 36.2 to 610.1 ± 124.4 mm³; p < 0.0001). In contrast, RN1734-treated rats showed no progression of ventricular enlargement, with mean ventricular volume decreasing from 137.2 ± 25.0 mm³ at P7 to 109.9 ± 35.5 mm³ at P15. Similarly, ventricular volume remained stable following triple combination therapy (163.6 ± 23.5 and 165.3 ± 40.2 mm³ at P7 and P15, respectively; p = 0.9744; Fig. 2i). These findings suggest that TRPV4 antagonism or triple combination therapy stabilizes ventricular volume near baseline levels and prevents progression of ventriculomegaly.

The change (Δ) in ventricular volume from P7 to P15 differed significantly among treatment groups in *Tmem*67□□□ animals (Fig. 2j). Δ volume did not differ significantly between untreated and EPO+MLT-treated animals (360.9 ± 59.7 mm³ vs. 407.1 ± 96.7 mm³; p = 0.9933), both of which showed significantly greater Δ volume than animals treated with RN alone (1.7 ± 40.1 mm³; both p < 0.0001) or with triple combination therapy (3.2 ± 29.3 mm³; both p < 0.0001). Δ volume did not differ significantly between RN-treated and triple-combination-treated animals (p > 0.9999). No significant differences in Δ volume were observed among treatment groups in *Tmem*67□□□ animals (Normal; all p > 0.9999). Together, these differences in Δ ventricular volume across genotypes and treatment groups further support the contention that the TRPV4 antagonist and triple combination therapy both mitigate progression of ventriculomegaly in homozygous (*Tmem*67□□□) animals.

### Treatment response in the *Tmem*67□□□ animals correlated with baseline ventricular severity

To provide a more detailed view of ventricular growth and the effect of treatment, the comparisons of individual animals in all cohorts are shown in Figure 3. As illustrated in the composite data, in the outbred Wistar rats, the ventricles showed a normal growth pattern which was unaffected by drug treatment with one exception (one animal in the triple cotreatment cohort). The same pattern is true in the *Tmem*67□□□ where the starting ventricle volumes are higher at P7 and the growth of the ventricles is, on average, more robust (Fig. 3b). All the untreated animals with hydrocephalus have higher ventricular volumes at P7 and this trend increases dramatically by P15 (Fig. 3c).

**Fig. 3.**
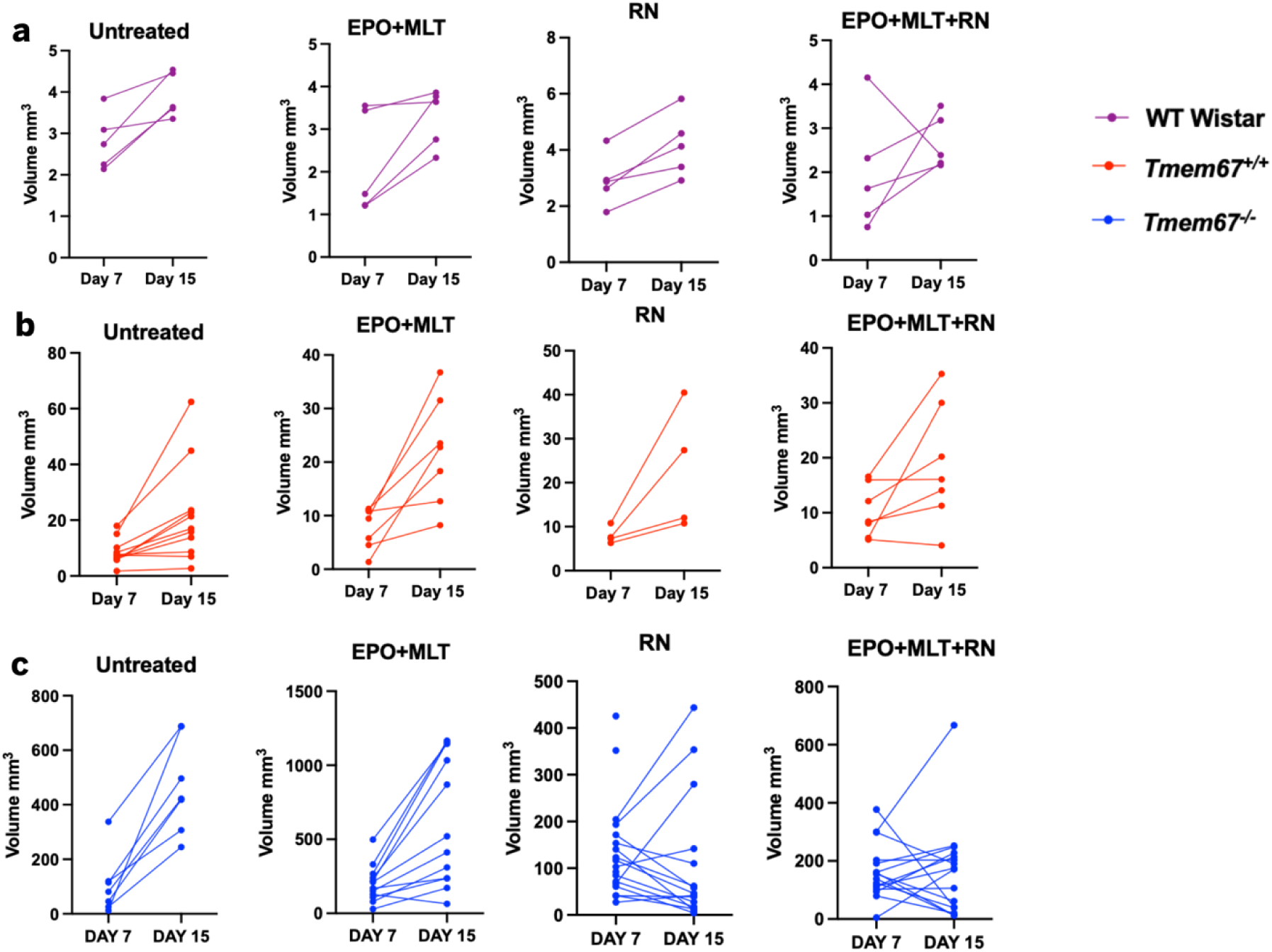
Individual animal trajectories showing progression of ventriculomegaly from Day 7 to Day 15 across strains and treatment groups. (a–c) Ventricular volume for individual animals at Day 7 and Day 15, connected by lines to show each animal’s trajectory over time, in Control Wistar (a), *Tmem*67□□□ (b), and *Tmem*67□□□ (c) rats. Within each row, animals are grouped by treatment conditions: untreated, EPO+MLT, RN1734, or EPO+MLT+RN1734. Two animals lack a Day 15 value because they did not survive to P15 (c, RN treated). n≥4 per group. These data are provided for illustrative comparison; no statistical comparisons were performed.

Among treated *Tmem*67□□□ animals more broadly, individual trajectories were variable: some animals showed volume increases from P7 to P15, while others showed relatively stable or decreased volume. This variability appeared related to baseline (P7) volume, with animals showing lower P7 volume tending to show more stable or decreasing trajectories, and animals with higher P7 volume tending to show continued expansion, consistent with the range of expansion illustrated in Fig. 2a–e. Only one of the hydrocephalic animals treated with EPO+MLT showed a decrease from P7 to P15 while approximately half of the TRPV4 agonist or triple co-therapy animals showed ventricular decreases over the treatment window.

### Hydrocephalus is associated with structural disruption of the periventricular white matter and ependymal lining

To characterize hydrocephalus-associated structural alterations within the periventricular region, H&E-stained coronal brain sections were examined at comparable rostrocaudal levels across untreated *Tmem*67□□□, untreated *Tmem*67□□□, and *Tmem*67□□□ treated with EPO+MLT, RN1734, and EPO+MLT+RN1734 (Fig. 4). The schematic in Fig. 4a illustrates the rostrocaudal sampling strategy and representative coronal level selected for histological assessment.

**Fig. 4.**
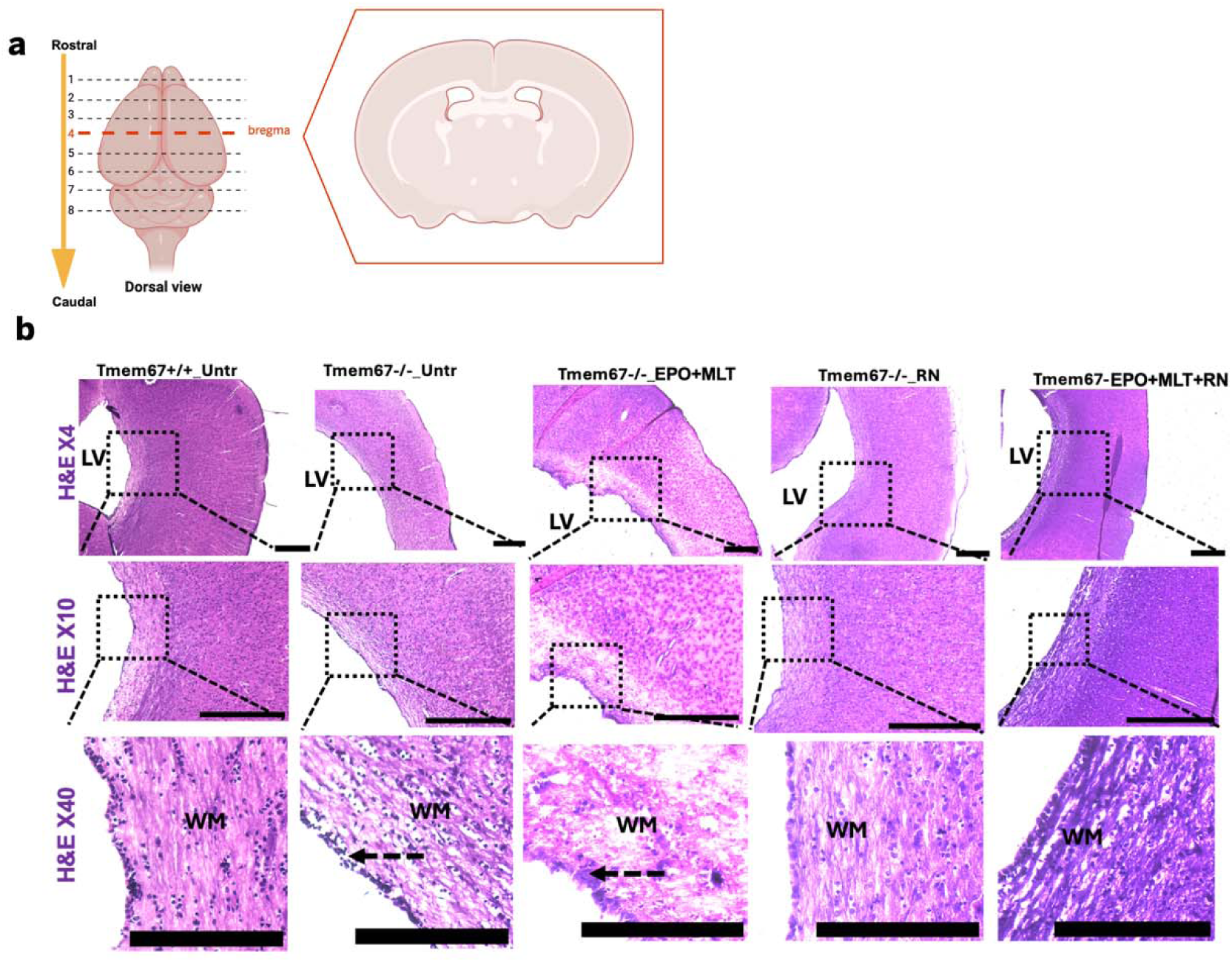
Hydrocephalus-associated structural alterations in the periventricular white matter and ependymal region. (a) Schematic illustrating the rostrocaudal orientation and coronal brain level used for histological analysis. The dashed lines indicate representative rostrocaudal levels, with the selected coronal section corresponding to the region containing the lateral ventricles and surrounding periventricular tissue. Created with BioRender.com. (b) Representative H&E-stained coronal brain sections from untreated *Tmem*67□□□, untreated *Tmem*67□□□, EPO+MLT-treated *Tmem*67□□□, RN1734-treated *Tmem*67□□□, and EPO+MLT+RN1734-treated *Tmem*67□□□ rats. Scale bars: 100 µm and high magnification images show the lateral ventricle (LV) and adjacent periventricular white matter (WM). Untreated *Tmem*67□□□ rats exhibited ventricular enlargement and structural alterations within the adjacent periventricular tissue, including disruption of the ependymal lining (black dashed arrow) Scale bar: 15 µm. Treatment groups showed variable preservation of periventricular tissue architecture, with comparatively more organized white-matter morphology observed in the RN1734– and EPO+MLT+RN1734-treated groups.

At low magnification (×4 and ×10), untreated *Tmem*67□□□ controls exhibited preserved periventricular tissue architecture surrounding the lateral ventricle (LV). In contrast, untreated *Tmem*67□□□ rats showed marked ventricular enlargement accompanied by distortion and compression of the adjacent periventricular tissue (Fig. 4b). Higher-magnification (×40) examination further revealed prominent structural alterations within the periventricular white matter (WM) of untreated *Tmem*67□□□ and EPO+MLT-treated *Tmem*67□□□ rats, including disrupted ependymal lining (EL) (black dashed arrow; Fig. 4b). Treatment with RN1734, or EPO+MLT+RN1734 was associated with variable preservation of periventricular tissue architecture compared with untreated *Tmem*67□□□ rats and EPO+MLT. (Fig. 4b).

### EPO+MLT, RN1734, and triple combination therapy attenuate elevated TLR4 immunoreactivity in the hydrocephalic rat periventricular region

To determine whether hydrocephalus-associated structural injury within the periventricular WM and EL was accompanied by enhanced innate inflammatory signaling potentially associated with damage-associated molecular patterns (DAMP) and to assess the effects of treatment on this response, TLR4 expression was examined in the periventricular region of untreated *Tmem*67□□□, untreated *Tmem*67□□□, and *Tmem*67□□□ rats treated with EPO+MLT, RN1734, or EPO+MLT+RN1734 (Fig. 5a-d). Representative immunofluorescence images revealed relatively low TLR4 immunoreactivity in untreated *Tmem*67□□□ controls, whereas untreated *Tmem*67□□□ rats exhibited markedly increased TLR4 signal within the periventricular region, including the WM and EL (white dashed arrows; Fig. 5a).

**Fig. 5.**
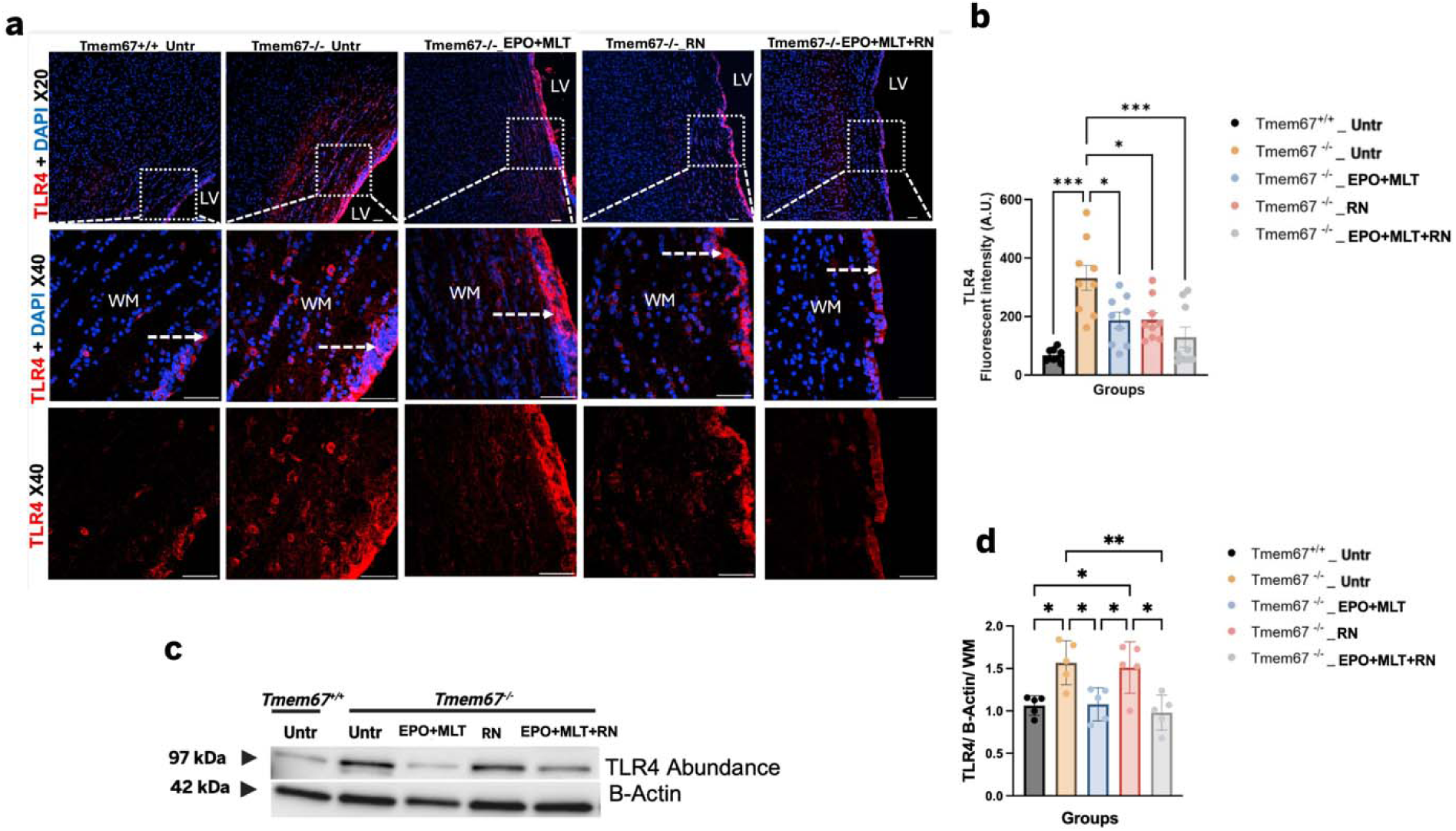
EPO+MLT, RN1734, and triple combination therapy attenuate elevated TLR4 immunoreactivity in the hydrocephalic rat white matter. (a) Representative immunofluorescence images showing TLR4 (red) and DAPI (blue) in the periventricular region of untreated *Tmem67*□□□, untreated *Tmem67*□□□, EPO+MLT-treated *Tmem*67□□□, RN1734-treated *Tmem*67□□□, and EPO+MLT+RN1734-treated *Tmem*67□□□ rats. Scale bars: 50 µm (low magnification) and 15 µm (high magnification). White dashed boxes indicate regions shown at higher magnification, and white dashed arrows indicate the ependymal lining adjacent to the lateral ventricle (LV). WM indicates the adjacent white matter. (b) Quantification of TLR4 fluorescence intensity within the periventricular white matter demonstrated significantly increased TLR4 immunoreactivity in untreated *Tmem*67□□□ rats compared with *Tmem*67□□□ controls. EPO+MLT, RN1734, and EPO+MLT+RN1734 treatments significantly reduced TLR4 fluorescence intensity relative to untreated *Tmem*67□□□ rats. Data are presented as mean ± Standard Error of the Mean (SEM) (n = 3 litters; 3 brain sections per animal). *p < 0.05, **p < 0.01, ***p < 0.001 (c) Representative Western blots, TLR4 (∼97 kDa), and β-Actin (∼42 kDa, loading control) in white matter (WM) from *Tmem67*□□□ untreated animals and *Tmem67*□□□ animals that were untreated or treated with EPO+MLT, RN1734, or EPO+MLT+RN1734. (d) Western blot quantification TLR4 band intensity, normalized to β-Actin in WM. *Tmem67*□□□ untreated animals are shown as a healthy baseline comparator; treatment groups (EPO+MLT, RN1734, EPO+MLT+RN1734) were tested only in *Tmem*67□□□ animals. Statistical significance was determined by one-way ANOVA with Tukey’s multiple comparisons test. Data are presented as mean ± SD, n≥4 per group. *p<0.05, **p<0.01, ***p<0.001, ****p<0.0001; ns, not significant. Data are presented as mean ± SEM (*Tmem67*□□□ Untr: n=5; *Tmem67*□□□ Untr: n=5; *Tmem67*□□□ EPO+MLT: n=5; *Tmem67*□□□ RN: n=5; *Tmem67*□□□ EPO+MLT+RN: n=5). *p<0.05, **p<0.01, ***p<0.001, ****p<0.0001

Quantitative fluorescence analysis confirmed a significant increase in TLR4 immunoreactivity within the WM of untreated *Tmem*67□□□ rats (332.8 ± 42.69 A.U.) compared with untreated *Tmem*67□□□ controls (68.58 ± 7.00 A.U.; p < 0.001; Fig. 5b). EPO+MLT, RN1734, and EPO+MLT+RN1734 treatments significantly reduced TLR4 fluorescence intensity to 188.1 ± 27.65 A.U. (p < 0.05), 189.2 ± 24.07 A.U. (p < 0.05), and 130.2 ± 34.70 A.U. (p < 0.001), respectively, compared with untreated *Tmem*67□□□ rats (Fig. 5b). These findings demonstrated a treatment responsive increase in TLR4 immunoreactivity localized to the structurally affected periventricular region.

Western blot analysis further supported the increased TLR4 protein abundance in hydrocephalic WM tissue, with TLR4 protein abundance (normalized to β-actin) significantly elevated in untreated *Tmem*67□□□ rats (1.568 ± 0.116) compared with untreated *Tmem*67□□□ controls (1.063 ± 0.052; p = 0.0155; Fig. 5c, d). EPO+MLT and EPO+MLT+RN1734 treatment significantly reduced TLR4 abundance to 1.078 ± 0.087 (p = 0.0193) and 0.981 ± 0.092 (p = 0.0043), respectively, compared with untreated *Tmem*67□□□ rats, whereas RN1734 alone did not significantly reduce TLR4 abundance (1.510 ± 0.136; p = 0.9940; Fig. 5d). To determine whether the observed TLR4 alterations extended beyond the periventricular WM, TLR4 protein abundance was additionally examined in the cortex (CTX), subventricular zone (SVZ), and choroid plexus (CPX) (Supplementary Fig. 1b). Regional Western blot analysis demonstrated detectable TLR4 expression in all three regions; however, no significant differences in TLR4 abundance were observed among experimental groups within the CTX, SVZ, or CPX (Supplementary Fig. 1a–e). These findings suggest that the pronounced treatment-responsive TLR4 changes observed in the present study are regionally associated with the structurally affected periventricular WM rather than reflecting a uniform alteration in TLR4 abundance throughout the brain and CPX.

### RN1734 and triple combination therapy attenuate regionally elevated NLRP3 expression in the hydrocephalic periventricular region

To determine whether treatment-associated attenuation of TLR4 within the periventricular WM was accompanied by changes in NLRP3-associated inflammatory signaling, NLRP3 expression was examined in untreated *Tmem*67□□□, untreated *Tmem*67□□□, and *Tmem*67□□□ rats treated with EPO+MLT, RN1734, or triple combination therapy (Fig. 6a–d). Representative immunofluorescence images demonstrated relatively low NLRP3 immunoreactivity in untreated *Tmem*67□□□ controls, whereas untreated *Tmem*67□□□ rats exhibited markedly increased NLRP3 signal within the periventricular region, including the WM and EL.

**Fig. 6.**
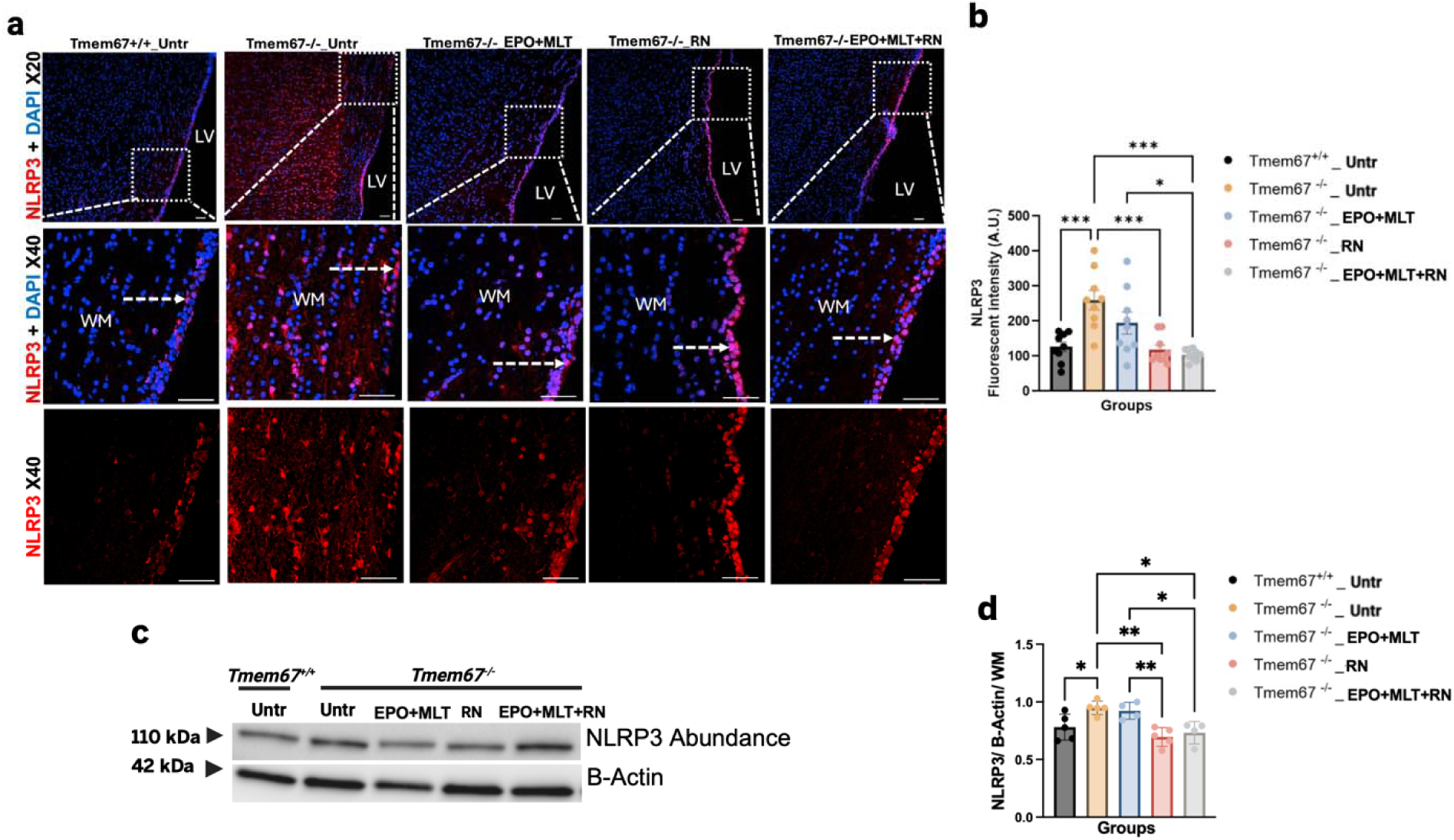
RN1734 and triple combination therapy attenuate elevated NLRP3 expression in the periventricular region of hydrocephalic rats. (a) Representative immunofluorescence images showing NLRP3 (red) and DAPI (blue) in the periventricular region of untreated *Tmem*67□□□, untreated *Tmem*67□□□, EPO+MLT-treated *Tmem*67□□□, RN1734-treated *Tmem*67□□□, and EPO+MLT+RN1734-treated *Tmem*67□□□ rats. Scale bars: 50 µm (low magnification) and 15 µm (high magnification). White dashed boxes indicate regions shown at higher magnification, and white dashed arrows indicate the ependymal lining adjacent to the lateral ventricle (LV). WM indicates the adjacent white matter. (b) Quantification of NLRP3 fluorescence intensity demonstrated significantly increased NLRP3 immunoreactivity in untreated *Tmem*67□□□ rats compared with *Tmem*67□□□ controls. EPO+MLT, RN1734, and EPO+MLT+RN1734 treatments significantly reduced NLRP3 fluorescence intensity compared with untreated *Tmem*67□□□ rats. Data are presented as mean ± Standard Error of the Mean (SEM) (n = 3 litters; 3 brain sections per animal). *p < 0.05, **p < 0.01, ***p < 0.001 (c) Representative Western blots, NLRP3 (∼110 kDa), and β-Actin (∼42 kDa, loading control) in white matter (WM) from *Tmem*67□□□ untreated animals and *Tmem*67□□□ animals that were untreated or treated with EPO+MLT, RN1734, or EPO+MLT+RN1734. (d) Western blot quantification of NLRP3 band intensity, normalized to β-Actin, in WM. *Tmem*67□□□ untreated animals are shown as a healthy baseline comparator; treatment groups (EPO+MLT, RN1734, EPO+MLT+RN1734) were tested only in *Tmem*67□□□ animals. Statistical significance was determined by one-way ANOVA with Tukey’s multiple comparisons test. Data are presented as mean ± SEM (*Tmem*67□□□ Untr: n=5; *Tmem*67□□□ Untr: n=5; *Tmem*67□□□ EPO+MLT: n=4; *Tmem*67□□□ RN: n=5; *Tmem*67□□□ EPO+MLT+RN: n=4). *p<0.05, **p<0.01, ***p<0.001, ***p<0.0001; ns, not significant.

Quantitative fluorescence analysis confirmed a significant increase in NLRP3 immunoreactivity within the periventricular WM of untreated *Tmem*67□□□ rats (260.0 ± 27.96 A.U.) compared with untreated *Tmem*67□□□ controls (126.2 ± 13.91 A.U.; p < 0.001; Fig. 6b). RN1734, and EPO+MLT+RN1734 treatments significantly reduced NLRP3 fluorescence intensity to 117.7 ± 13.53 A.U. (p < 0.05), and 103.5 ± 5.52 A.U. (p < 0.001), respectively, compared with untreated *Tmem*67□□□ rats (Fig. 6b). These findings demonstrate increased NLRP3 immunoreactivity within the structurally affected periventricular region and its attenuation following treatment.

Western blot analysis further demonstrated increased NLRP3 protein abundance within the periventricular WM, with NLRP3 protein abundance (normalized to β-actin) significantly elevated in untreated *Tmem*67□□□ rats (0.949 ± 0.026) compared with untreated *Tmem*67□□□ controls (0.780 ± 0.051; p = 0.0466; Fig. 6c, d). RN1734 and EPO+MLT+RN1734 treatment significantly reduced NLRP3 abundance to 0.696 ± 0.036 (p = 0.0018) and 0.732 ± 0.049 (p = 0.0118), respectively, compared with untreated *Tmem*67□□□ rats, whereas EPO+MLT alone did not significantly reduce NLRP3 abundance (0.922 ± 0.037; p = 0.9909; Fig. 6d).

To determine whether these NLRP3 alterations extended beyond the periventricular WM, NLRP3 protein abundance was additionally examined in the CTX, SVZ, and CPX (Supplementary Fig. 2b). NLRP3 was detectable in all three regions; however, no significant differences in NLRP3 protein abundance were observed among experimental groups within the CTX, SVZ, or CPX (Supplementary Fig. 2b–e). Together, these results indicate that the pronounced treatment-responsive changes in NLRP3 observed in the present study were localized predominantly to the structurally affected periventricular WM rather than representing a uniform alteration in NLRP3 abundance across all examined brain regions.

### TRPV4-stimulated increase in transepithelial electrolyte transport and barrier permeability is unaffected by short– or long-term EPO+MLT treatment

Given the role of the choroid plexus epithelium in maintaining CSF homeostasis, changes in epithelial ion transport and barrier permeability could contribute to altered CSF dynamics and ventricular enlargement. As TRPV4 is predominantly expressed on the apical side of the choroid plexus epithelium and is involved in regulating ion transport and permeability, we examined whether EPO+MLT treatment alters TRPV4-dependent epithelial responses and whether this could be related to the changes in ventricular size observed in the rat model, using a well-characterized human choroid plexus epithelial cell model (HIBCPP) [19, 22].

To assess TRPV4-dependent changes in epithelial electrolyte flux and barrier function, we measured short-circuit current (Isc; a measure of net electrogenic ion flux) and transepithelial conductance (a measure of transepithelial permeability) with Ussing chamber electrophysiological techniques before and after activation of TRPV4 with GSK1016790A in HIBCPP cells. Baseline Isc and conductance were unaffected by EPO+MLT treatment alone, and GSK1016790A produced a comparable biphasic Isc response and a comparable progressive rise in conductance in EPO+MLT-treated and control groups following both short-term (10 min; Fig. 7a, b) and long-term (24 h; Fig. 7c, d) exposure. Together, these data indicate that TRPV4-mediated epithelial electrolyte flux and barrier permeability are not affected by EPO+MLT treatment, regardless of treatment duration.

**Fig. 7.**
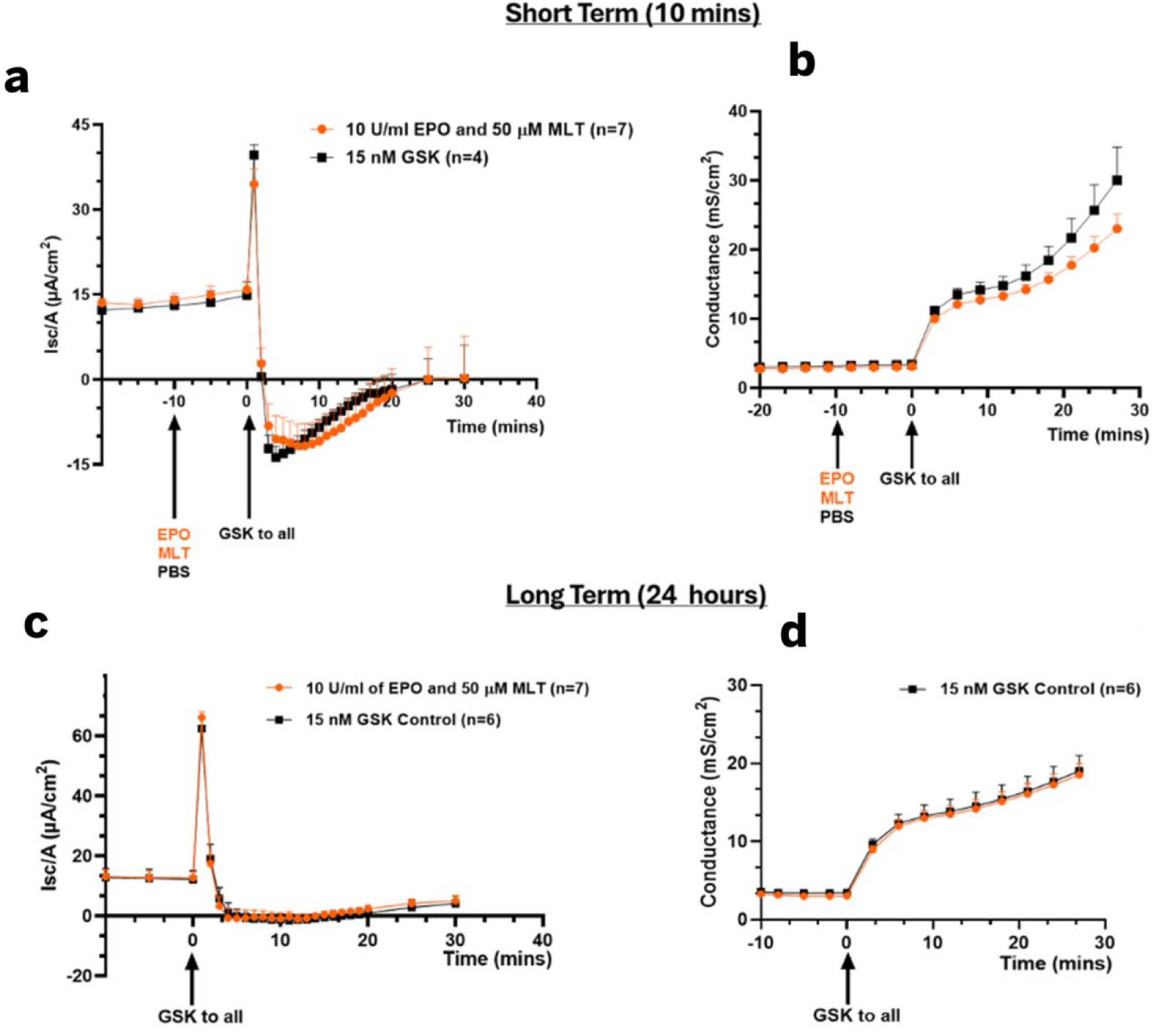
EPO+MLT treatment does not affect net transepithelial ion flux or membrane permeability in a human choroid plexus epithelial cell line HIBCPP. Ussing-style electrophysiology was used to determine: (a and c) TRPV4-stimulated (GSK1016790A; 15 nM) changes in the short-circuit current (Isc; a measure of net transepithelial electrolyte flux); and (b and d) changes in membrane conductance (mS/cm^2^ a measure of transepithelial permeability) following short-term (10 min; a and b) or long term (24 hours; c and d) treatment with EPO and MLT applied to both sides of the confluent monolayers of the human choroid plexus cell line HIBCPP. The number of individual monolayers is indicated in the figure legends. Statistical analyses were performed using multiple Student’s t-tests. No statistical differences were observed with EPO+MLT treatment.

## Discussion

Previous studies established that the *Tmem*67□□□ rat develops progressive ventriculomegaly accompanied by increased TLR4– and NLRP3-associated inflammatory signaling and demonstrated that TRPV4 antagonism with RN1734 attenuates ventricular enlargement and inflammatory pathology [3, 18]. Building on these findings, the present study addresses a different therapeutic question: whether combining RN1734 with the neuroprotective, anti-inflammatory agents EPO and MLT could provide broader protection against the multiple pathological consequences of congenital hydrocephalus. We therefore compared EPO+MLT, RN1734, and triple combination therapy across cranial, ventricular, periventricular structural, and inflammatory outcomes. Our findings demonstrate that the triple combination therapy attenuated ventricular progression (Fig. 2) and cranial enlargement (Fig. 1b, c) while also reducing TLR4– (Fig. 5) and NLRP3-associated (Fig. 6) inflammatory signaling within the structurally affected periventricular white matter. These findings extend our previous work by evaluating a multi-target therapeutic strategy designed to simultaneously address CSF/ventricular pathology and secondary parenchymal injury and inflammation.

Consistent with our previous studies, untreated *Tmem*67□□□ rats exhibited rapid progression of ventriculomegaly between P7 and P15, and RN1734 markedly attenuated this progression (Fig. 2i)[18]. EPO+MLT treatment alone did not prevent ventricular expansion in our genetic rat model, whereas the EPO+MLT+RN1734 combination maintained ventricular volume near pretreatment levels (Fig. 2h, i). Analysis of the change in ventricular volume demonstrated no significant difference between RN1734 and triple-combination treatment (Fig. 2j), indicating that RN1734 is likely the principal contributor to the ventricular-stabilizing effect of the combination in this injury model. Thus, the therapeutic rationale for adding EPO+MLT is not necessarily to further enhance the effect of TRPV4 antagonism on ventricular expansion in *Tmem*67□□□ rats, but rather to address additional pathological consequences of hydrocephalus that may not be adequately targeted through modulation of CSF dynamics alone [39, 49].

Cranial measurements provided an additional anatomical measure of hydrocephalus progression during development [18, 41]. Untreated and EPO+MLT-treated *Tmem*67□□□ rats maintained significantly greater cranial dimensions than their corresponding *Tmem*67□□□ controls (Fig. 1b, c). In contrast, the genotype-associated difference was no longer significant following RN1734 treatment while, interestingly, triple-combination-treated *Tmem*67□□□ animals exhibited significantly smaller vertical and horizontal cranial dimensions than similarly treated *Tmem*67□□□ animals (Fig. 1b, c). These observations suggest that treatments containing RN1734 substantially modify macrocephaly with progressive hydrocephalus in skulls that are not ossified. However, cranial dimensions are also influenced by overall postnatal head and body growth, and *Tmem*67□□□ animals remained significantly lighter than *Tmem*67□□□ littermates across treatment conditions (Fig. 1d). Cranial measurements should therefore be interpreted together with body weight measurements, postnatal age, and MRI-derived ventricular volume in combination rather than as an independent measure of ventricular pathology at this developmental stage.

The identification of periventricular WM and ependymal abnormalities provided the rationale for examining innate inflammatory signaling within these structurally affected regions. The elevation of TLR4 and NLRP3 immunoreactivity in the periventricular WM and ependymal of untreated *Tmem*67□□□ rats is consistent with a DAMP-associated inflammatory response accompanying ventricular enlargement and ependymal disruption [34, 45]. All three treatment regimens reduced these markers to varying degrees, with EPO+MLT and the triple combination most effective in mitigating the increase in TLR4, and RN1734 and the triple combination most effective against NLRP3. Notably, the triple combination significantly reduced both TLR4 and NLRP3, suggesting broader attenuation of ubiquitous inflammatory signaling. Together with the comparatively preserved periventricular tissue architecture observed following treatment, these findings suggest that triple combination therapy may address both the structural and inflammatory consequences of progressive hydrocephalus. It is important to also note that the absence of significant treatment associated differences in TLR4 and NLRP3 abundance within the CTX, SVZ, and CPX (Supplementary Fig.1b-e and Fig. 2b-e), despite marked changes in periventricular WM (Fig. 5c, d and Fig. 6 c, d) and EL (Supplementary Fig. 1a and 2a) suggests that TLR4-associated inflammatory signaling is spatially heterogeneous in genetic hydrocephalus pathophysiology. The localization of increased TLR4 and NLRP3 to regions exhibiting prominent structural injury (WM and EL) (Fig. 4b) supports an association between local tissue disruption and inflammation [34]. Notably, EPO+MLT treatment did not alter TRPV4-mediated short-circuit current or conductance responses in human choroid plexus epithelial cells (HIBCPP), regardless of the treatment duration. This finding indicates that the beneficial effects of EPO+MLT observed *in vivo* are not mediated through modulation of TRPV4 channel activity at the choroid plexus epithelium (Fig. 7).

The principal novelty of the present study lies in the multi-target therapeutic strategy. Congenital hydrocephalus is not solely a disorder of excessive ventricular CSF accumulation [3, 17, 47, 49]. Progressive ventriculomegaly in the neonatal period is accompanied by cranial enlargement, mechanical deformation of surrounding brain tissue, ependymal and white-matter injury, inflammatory signaling, oxidative stress, and cellular damage [12, 32]. A treatment directed primarily at CSF/ventricular pathology may therefore not address all secondary consequences of the disease, whereas a neuroprotective or anti-inflammatory intervention that does not control ventricular expansion may likewise provide incomplete protection [47, 49]. Our results revealed a mechanistically targeted triple therapy cocktail yielded a broad treatment response encompassing cranial dimensions, periventricular tissue pathology, and both TLR4– and NLRP3-associated inflammatory signaling.

This study is constrained by scope and has limitations. Response to treatment among *Tmem67*□□□ animals was related to baseline (P7) ventricular severity, indicating that more severely affected animals may require adjusted treatment strategies. In addition, RN1734-treated and triple-combination-treated *Tmem*67□□□ animals showed lower body weight than untreated animals despite the anatomical benefits of treatment, an effect that will need to be characterized further to distinguish drug-related toxicity from disease-related growth impairment. Overall, these results support triple combination therapy with EPO, MLT, and a TRPV4 antagonist as a promising non-invasive pharmacological strategy for congenital hydrocephalus, warranting further studies to define optimal dosing, long-term safety, and the precise mechanistic contribution of each agent.

## Conclusion

In conclusion, the present study demonstrates that triple combination therapy with EPO, MLT, and a TRPV4 antagonist targets multiple pathological features of congenital hydrocephalus in *Tmem*67□□□^−/−^ rats. Triple combination therapy attenuated ventricular progression and cranial enlargement, while reducing TLR4– and NLRP3-associated inflammatory signaling within the structurally affected periventricular WM region. These findings support a multi-target therapeutic approach for the treatment of congenital hydrocephalus (Fig. 8).

**Fig. 8.**
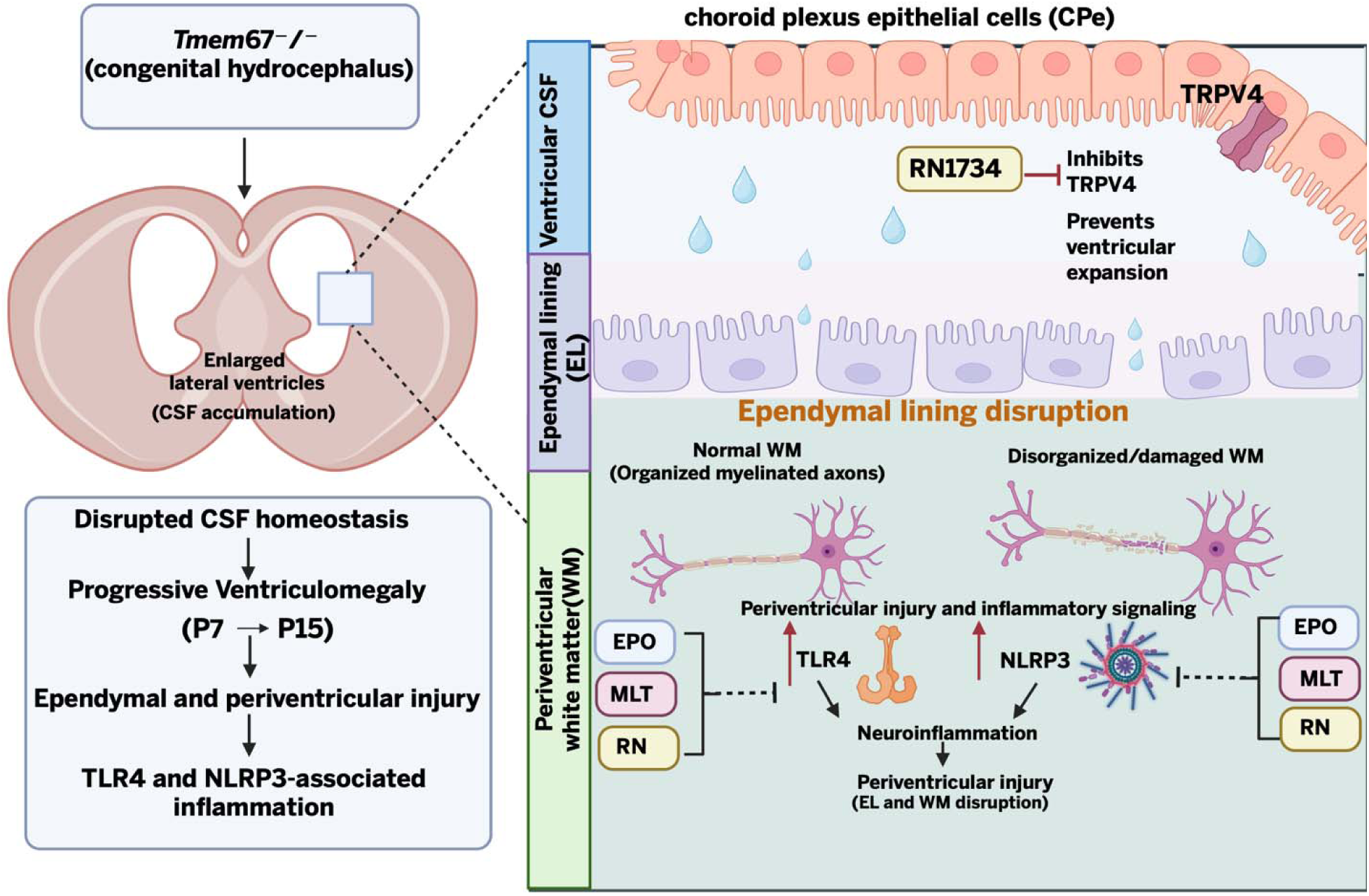
Proposed mechanisms of ependymal and periventricular white matter injury in Tmem67□□□ congenital hydrocephalus and potential therapeutic targets. (Left, top) Loss of *Tmem67* in the *Tmem67*□□□ rat causes congenital hydrocephalus. The schematic coronal brain section shows enlarged lateral ventricles resulting from CSF accumulation. (Left, bottom) Proposed sequence of disease progression: disrupted CSF homeostasis drives progressive ventriculomegaly between postnatal day 7 (P7) and P15 leading to ependymal and periventricular injury, followed by TLR4– and NLRP3 inflammasome-associated inflammation. (Right) Magnified view of the boxed periventricular region showing the ventricular CSF, the ependymal lining (EL), and the underlying periventricular white matter (WM). Choroid plexus epithelial cells (CPe) express the TRPV4 channel and contribute to CSF accumulation within the ventricle. The TRPV4 antagonist RN1734 (RN) inhibits TRPV4 and prevents ventricular expansion. Ventricular enlargement is accompanied by disruption of the EL, with loss of ependymal cell continuity allowing fluid to move into the periventricular tissue. Within the periventricular WM, normal tissue with organized myelinated axons becomes disorganized and damaged, with myelin fragmentation. This periventricular injury is associated with upregulation (red arrows) of TLR4 and NLRP3 inflammasome signaling, which promotes neuroinflammation and further periventricular injury affecting both the EL and WM. Combination therapy with EPO, MLT, and RN is proposed to attenuate TLR4– and NLRP3-mediated inflammatory signaling, thereby limiting neuroinflammation and secondary injury. Black arrows indicate progression or downstream effects. Solid blunt-ended lines indicate inhibition and dashed blunt-ended lines indicate proposed inhibition. Created with BioRender.com.

## Acknowledgements

The authors thank Cameryn Riley Davis for her assistance with the preparation and maintenance of cell cultures used in this study, and Kelsey Stabler for her assistance with MRI analysis.

## Funding Declaration

This work was supported by The Assistant Secretary of Defense for Health Affairs endorsed by the Department of Defense through the Peer Reviewed Medical Research Program under Award Number (HT9425-23-1-0296).

Opinions, interpretations, conclusions, and recommendations are those of the author and are not necessarily endorsed by The Assistant Secretary of Defense for Health Affairs endorsed by the Department of Defense.

## Contribution

F.O.B, B.B.Y wrote the main manuscript text, F.O.B prepared figures, F.O.B, D.A., G. M., Y.Z. conducted experiments, F.O.B, D.A., G. M. and B.B.Y. analyzed data, L.L.J., S.R. provided feedback on early drafts of data and manuscript, all authors read and approved the final manuscript.

## Ethics and Consent to Participate declarations

not applicable.

## References

1. Abdelhamed ZA, Natarajan S, Wheway G, Inglehearn CF, Toomes C, Johnson CA, Jagger DJ (2015) The Meckel-Gruber syndrome protein TMEM67 controls basal body positioning and epithelial branching morphogenesis in mice via the non-canonical Wnt pathway. Dis Model Mech 8:527–41. 10.1242/dmm.019083

2. Anwar F, Zhang K, Sun C, Pang M, Zhou W, Li H, et al. (2024) Hydrocephalus: An update on latest progress in pathophysiological and therapeutic research. Biomed Pharmacother 181:117702. 10.1016/j.biopha.2024.117702

3. Audu D, Reed M, Blazer-Yost BL (2026) TRPV4 antagonism attenuates TLR4-NLRP3-mediated neuroinflammation and oxidative stress in cingulate cortex neurons in a genetic rat model of hydrocephalus. J Neuroinflammation 10.1186/s12974-026-04031-7

4. Carrillo-Vico A, Lardone PJ, Alvarez-Sánchez N, Rodríguez-Rodríguez A, Guerrero JM (2013) Melatonin: buffering the immune system. Int J Mol Sci 14:8638–83. 10.3390/ijms14048638

5. Cho YK, Kim G, Park S, Sim JH, Won YJ, Hwang CH, et al. (2012) Erythropoietin promotes oligodendrogenesis and myelin repair following lysolecithin-induced injury in spinal cord slice culture. Biochem Biophys Res Commun 417:753–9. 10.1016/j.bbrc.2011.12.029

6. Constantinescu C, Constantinescu Ziegelitz D, Wikkelsö C, Kern S, Jaraj D, Westman E, et al. (2025) Idiopathic Normal Pressure Hydrocephalus: Incidence, Prevalence, and Progression in a Large Population-Based Cohort of Older Adults. Neurology Open Access 1:e000035. 10.1212/WN9.0000000000000035

7. Deng Z, Wang H, Zhong K, Li Y, Deng H, Gao B, et al. (2025) The Role of Choroid Plexus in Hydrocephalus from the Perspective of Structure and Function: a Therapeutic Target. Mol Neurobiol 62:9133–50. 10.1007/s12035-025-04823-7

8. Dewan MC, Rattani A, Mekary R, Glancz LJ, Yunusa I, Baticulon RE, et al. (2019) Global hydrocephalus epidemiology and incidence: systematic review and meta-analysis. J Neurosurg 130:1065–79. 10.3171/2017.10.Jns17439

9. Digicaylioglu M, Lipton SA (2001) Erythropoietin-mediated neuroprotection involves cross-talk between Jak2 and NF-kappaB signalling cascades. Nature 412:641–7. 10.1038/35088074

10. Drake JM, Kestle JR, Milner R, Cinalli G, Boop F, Piatt J, Jr., et al. (1998) Randomized trial of cerebrospinal fluid shunt valve design in pediatric hydrocephalus. Neurosurgery 43:294–303; discussion-5. 10.1097/00006123-199808000-00068

11. Farhadi N, Oryan S, Nabiuni M (2014) Serum levels of melatonin and cytokines in multiple sclerosis. Biomed J 37:90–2. 10.4103/2319-4170.125885

12. Garcia-Bonilla M, Castaneyra-Ruiz L, Zwick S, Talcott M, Otun A, Isaacs AM, et al. (2022) Acquired hydrocephalus is associated with neuroinflammation, progenitor loss, and cellular changes in the subventricular zone and periventricular white matter. Fluids Barriers CNS 19:17. 10.1186/s12987-022-00313-3

13. Gattone VH, 2nd, Tourkow BA, Trambaugh CM, Yu AC, Whelan S, Phillips CL, et al. (2004) Development of multiorgan pathology in the wpk rat model of polycystic kidney disease. Anat Rec A Discov Mol Cell Evol Biol 277:384–95. 10.1002/ar.a.20022

14. Gunata M, Parlakpinar H, Acet HA (2020) Melatonin: A review of its potential functions and effects on neurological diseases. Rev Neurol (Paris) 176:148–65. 10.1016/j.neurol.2019.07.025

15. Gunnarson E, Song Y, Kowalewski JM, Brismar H, Brines M, Cerami A, et al. (2009) Erythropoietin modulation of astrocyte water permeability as a component of neuroprotection. Proc Natl Acad Sci U S A 106:1602–7. 10.1073/pnas.0812708106

16. Hochstetler A, Hehnly C, Dawes W, Harris D, Sadegh C, Mangano FT, et al. (2025) Research priorities for non-invasive therapies to improve hydrocephalus outcomes. Fluids Barriers CNS 22:24. 10.1186/s12987-025-00632-1

17. Hochstetler A, Raskin J, Blazer-Yost BL (2022) Hydrocephalus: historical analysis and considerations for treatment. Eur J Med Res 27:168. 10.1186/s40001-022-00798-6

18. Hochstetler AE, Smith HM, Preston DC, Reed MM, Territo PR, Shim JW, et al. (2020) TRPV4 antagonists ameliorate ventriculomegaly in a rat model of hydrocephalus. JCI Insight 5:10.1172/jci.insight.137646

19. Hulme L, Hochstetler A, Schwerk C, Schroten H, Ishikawa H, Tung CY, et al. (2022) Characterization of TRPV4-mediated signaling pathways in an optimized human choroid plexus epithelial cell line. Am J Physiol Cell Physiol 323:C1823–c42. 10.1152/ajpcell.00193.2022

20. Imbesi M, Uz T, Manev H (2008) Role of melatonin receptors in the effects of melatonin on BDNF and neuroprotection in mouse cerebellar neurons. J Neural Transm (Vienna) 115:1495–9. 10.1007/s00702-008-0066-z

21. Isaacs AM, Yang R, Cadieux M, Ben-Israel D, Sader N, Opoku-Darko M, et al. (2023) Characteristics of shunt failure in 38,095 adult shunt insertion surgeries: a systematic review and meta-analysis. Neurosurg Focus 54:E2. 10.3171/2023.1.Focus22637

22. Ishiwata I, Ishiwata C, Ishiwata E, Sato Y, Kiguchi K, Tachibana T, et al. (2005) Establishment and characterization of a human malignant choroids plexus papilloma cell line (HIBCPP). Hum Cell 18:67–72. 10.1111/j.1749-0774.2005.tb00059.x

23. Iwai M, Stetler RA, Xing J, Hu X, Gao Y, Zhang W, et al. (2010) Enhanced oligodendrogenesis and recovery of neurological function by erythropoietin after neonatal hypoxic/ischemic brain injury. Stroke 41:1032–7. 10.1161/strokeaha.109.570325

24. Jantzie LL, Corbett CJ, Firl DJ, Robinson S (2015) Postnatal Erythropoietin Mitigates Impaired Cerebral Cortical Development Following Subplate Loss from Prenatal Hypoxia-Ischemia. Cereb Cortex 25:2683–95. 10.1093/cercor/bhu066

25. Jantzie LL, Miller RH, Robinson S (2013) Erythropoietin signaling promotes oligodendrocyte development following prenatal systemic hypoxic-ischemic brain injury. Pediatr Res 74:658–67. 10.1038/pr.2013.155

26. Juul SE, Anderson DK, Li Y, Christensen RD (1998) Erythropoietin and erythropoietin receptor in the developing human central nervous system. Pediatr Res 43:40–9. 10.1203/00006450-199801000-00007

27. Juul SE, Beyer RP, Bammler TK, McPherson RJ, Wilkerson J, Farin FM (2009) Microarray analysis of high-dose recombinant erythropoietin treatment of unilateral brain injury in neonatal mouse hippocampus. Pediatr Res 65:485–92. 10.1203/PDR.0b013e31819d90c8

28. Liu XY, Song X, Czosnyka M, Robba C, Czosnyka Z, Summers JL, et al. (2024) Congenital hydrocephalus: a review of recent advances in genetic etiology and molecular mechanisms. Mil Med Res 11:54. 10.1186/s40779-024-00560-5

29. Lutz BR, Venkataraman P, Browd SR (2013) New and improved ways to treat hydrocephalus: Pursuit of a smart shunt. Surg Neurol Int 4:S38–50. 10.4103/2152-7806.109197

30. Mahendran G, Torabi M, Blazer-Yost BL (2026) Phosphoproteomic Insights into TRPV4–AMPK Signaling Axis and Barrier Regulation in the Choroid Plexus Epithelium. International Journal of Molecular Sciences 27: https://www.mdpi.com/1422-0067/27/18/8305

31. Masuda S, Kada E, Nagao M, Sasaki R (1999) In vitro neuroprotective action of recombinant rat erythropoietin produced by astrocyte cell lines and comparative studies with erythropoietin produced by Chinese hamster ovary cells. Cytotechnology 29:207–13. 10.1023/a:1008014206833

32. McAllister JP, 2nd (2012) Pathophysiology of congenital and neonatal hydrocephalus. Semin Fetal Neonatal Med 17:285–94. 10.1016/j.siny.2012.06.004

33. Newland V, Jantzie LL, Blazer-Yost BL (2024) Understanding and modeling the pathophysiology of hydrocephalus: in search of better treatment options. Physiologia 10.3390/physiologia4020010

34. Pi W, Cao L, Wang Q, Yong VW, Yang L, Xue M (2026) The TLR4/MyD88 axis: orchestrator of neuroinflammation from molecular initiation to systemic crosstalk. Brain Hemorrhages 7:10.1016/j.hest.2026.03.002

35. Ramachandra R, Subramanian T. Atlas of the neonatal rat brain: CRC Press; 2016.

36. Rangarajan V, Juul SE (2014) Erythropoietin: emerging role of erythropoietin in neonatal neuroprotection. Pediatr Neurol 51:481–8. 10.1016/j.pediatrneurol.2014.06.008

37. Reiter RJ, Calvo JR, Karbownik M, Qi W, Tan DX (2000) Melatonin and its relation to the immune system and inflammation. Ann N Y Acad Sci 917:376–86. 10.1111/j.1749-6632.2000.tb05402.x

38. Rekate HL (2009) A contemporary definition and classification of hydrocephalus. Semin Pediatr Neurol 16:9–15. 10.1016/j.spen.2009.01.002

39. Robinson S, Conteh FS, Oppong AY, Yellowhair TR, Newville JC, Demerdash NE, et al. (2018) Extended Combined Neonatal Treatment With Erythropoietin Plus Melatonin Prevents Posthemorrhagic Hydrocephalus of Prematurity in Rats. Front Cell Neurosci 12:322. 10.3389/fncel.2018.00322

40. Sánchez-Barceló EJ, Mediavilla MD, Tan DX, Reiter RJ (2010) Clinical uses of melatonin: evaluation of human trials. Curr Med Chem 17:2070–95. 10.2174/092986710791233689

41. Shim JW, Territo PR, Simpson S, Watson JC, Jiang L, Riley AA, et al. (2019) Hydrocephalus in a rat model of Meckel Gruber syndrome with a TMEM67 mutation. Sci Rep 9:1069. 10.1038/s41598-018-37620-5

42. Shingo T, Sorokan ST, Shimazaki T, Weiss S (2001) Erythropoietin regulates the in vitro and in vivo production of neuronal progenitors by mammalian forebrain neural stem cells. J Neurosci 21:9733–43. 10.1523/jneurosci.21-24-09733.2001

43. Smith UM, Consugar M, Tee LJ, McKee BM, Maina EN, Whelan S, et al. (2006) The transmembrane protein meckelin (MKS3) is mutated in Meckel-Gruber syndrome and the wpk rat. Nat Genet 38:191–6. 10.1038/ng1713

44. Sugawa M, Sakurai Y, Ishikawa-Ieda Y, Suzuki H, Asou H (2002) Effects of erythropoietin on glial cell development; oligodendrocyte maturation and astrocyte proliferation. Neurosci Res 44:391–403. 10.1016/s0168-0102(02)00161-x

45. Teng W, Wang L, Xue W, Guan C (2009) Activation of TLR4-mediated NFkappaB signaling in hemorrhagic brain in rats. Mediators Inflamm 2009:473276. 10.1155/2009/473276

46. Tsai PT, Ohab JJ, Kertesz N, Groszer M, Matter C, Gao J, et al. (2006) A critical role of erythropoietin receptor in neurogenesis and post-stroke recovery. J Neurosci 26:1269–74. 10.1523/jneurosci.4480-05.2006

47. Wang H, Chen X, You C, Wu K, Sun T (2025) Navigating challenges in hydrocephalus following intraventricular hemorrhage: a comprehensive review of current evidence. Front Neurol 16:1630286. 10.3389/fneur.2025.1630286

48. Wang L, Zhang Z, Wang Y, Zhang R, Chopp M (2004) Treatment of stroke with erythropoietin enhances neurogenesis and angiogenesis and improves neurological function in rats. Stroke 35:1732–7. 10.1161/01.STR.0000132196.49028.a4

49. Zhao J, Tang Z (2024) Oxidative stress in hydrocephalus: a new potential therapeutic target. Brain-X 2:10.1002/brx2.70008

