## Supplementary file for "Therapeutic Potential of Melatonin, Erythropoietin, and TRPV4 Antagonism in a Preclinical Model of Genetic Hydrocephalus"

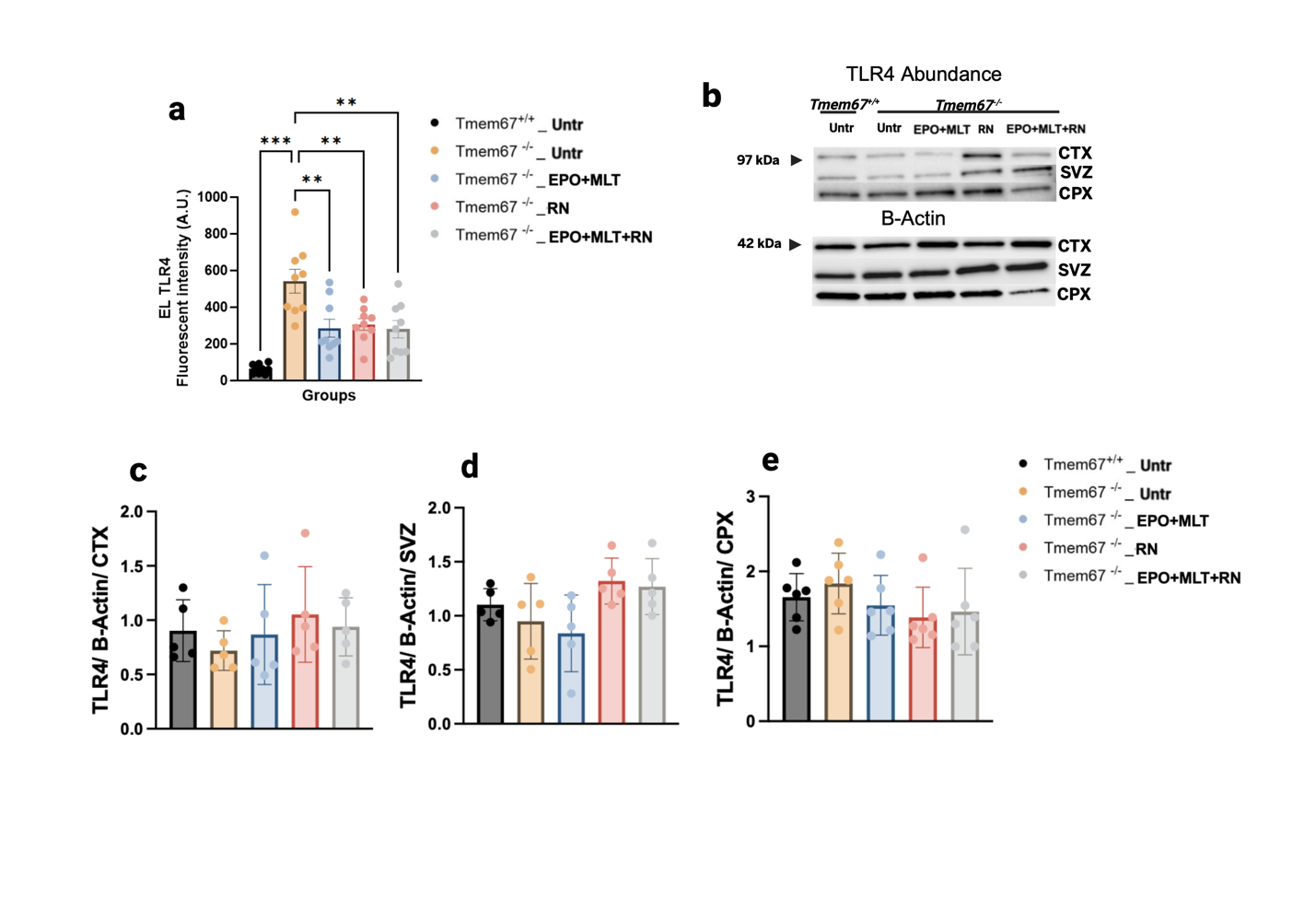

**Supplementary Fig. 1. TLR4 abundance across periventricular subregions in *Tmem*67⁻ᐟ⁻ rats**
(a) Quantification of TLR4 immunofluorescence intensity in the ependymal lining (EL) of untreated *Tmem*67⁺ᐟ⁺, untreated *Tmem*67⁻ᐟ⁻, and *Tmem*67⁻ᐟ⁻ rats treated with EPO+MLT, RN1734, or EPO+MLT+RN1734. Untreated *Tmem*67⁻ᐟ⁻ rats showed significantly increased EL TLR4 fluorescence intensity compared with *Tmem*67⁺ᐟ⁺ controls, and EPO+MLT, RN1734, and EPO+MLT+RN1734 treatments significantly reduced EL TLR4 fluorescence intensity relative to untreated *Tmem*67⁻ᐟ⁻ rats.
(b) Representative Western blots showing TLR4 (~97 kDa) and β-Actin (~42 kDa, loading control) abundance in cortex (CTX), subventricular zone (SVZ), and choroid plexus (CPX) from untreated *Tmem*67⁺ᐟ⁺ rats and *Tmem*67⁻ᐟ⁻ rats that were untreated or treated with EPO+MLT, RN1734, or EPO+MLT+RN1734.
(c–e) Quantification of TLR4 band intensity, normalized to β-Actin, in CTX (c), SVZ (d), and CPX (e). *Tmem*67⁺ᐟ⁺ untreated animals are shown as a healthy baseline comparator; treatment groups (EPO+MLT, RN1734, EPO+MLT+RN1734) were tested only in *Tmem*67⁻ᐟ⁻ animals. Statistical significance was determined by one-way ANOVA with Tukey's multiple comparisons test.
Data are presented as mean ± SEM (n = 3 litters; 3 brain sections per animal for a; n≥4 per group for c–e). *p<0.05, **p<0.01, ***p<0.001, ****p<0.0001; ns, not significant.
Abbreviations: EL, ependymal lining; CTX, cortex; SVZ, subventricular zone; CPX, choroid plexus; EPO, erythropoietin; MLT, melatonin; RN, RN1734.
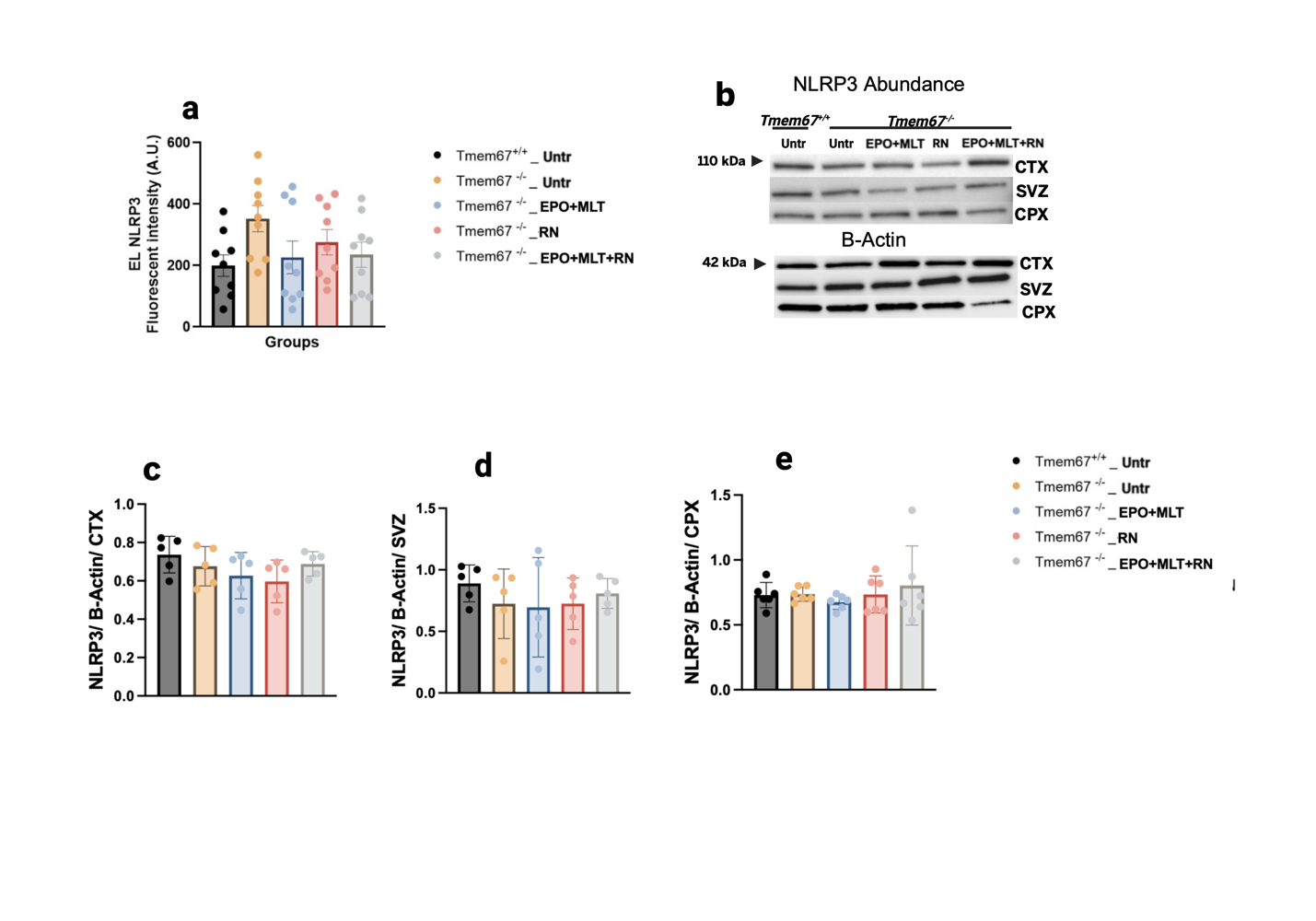

**Supplementary Fig. 2. NLRP3 abundance across periventricular subregions in *Tmem*67⁻ᐟ⁻ rats**
(a) Quantification of NLRP3 immunofluorescence intensity in the ependymal lining (EL) of untreated *Tmem*67⁺ᐟ⁺, untreated *Tmem*67⁻ᐟ⁻, and *Tmem*67⁻ᐟ⁻ rats treated with EPO/MLT, RN1734, or EPO+MLT+RN1734. No significant differences in EL NLRP3 fluorescence intensity were observed across groups.
(b) Representative Western blots showing NLRP3 (~110 kDa) and β-Actin (~42 kDa, loading control) abundance in cortex (CTX), subventricular zone (SVZ), and choroid plexus (CPX) from untreated *Tmem*67⁺ᐟ⁺ rats and *Tmem*67⁻ᐟ⁻ rats that were untreated or treated with EPO/MLT, RN1734, or EPO+MLT+RN1734.
(c–e) Quantification of NLRP3 band intensity, normalized to β-Actin, in CTX (c), SVZ (d), and CPX (e). *Tmem*67⁺ᐟ⁺ untreated animals are shown as a healthy baseline comparator; treatment groups (EPO/MLT, RN1734, EPO+MLT+RN1734) were tested only in *Tmem*67⁻ᐟ⁻ animals. Statistical significance was determined by one-way ANOVA with Tukey's multiple comparisons test.
Data are presented as mean ± SEM (n = 3 litters; 3 brain sections per animal for a; n≥4 per group for c–e). *p<0.05, **p<0.01, ***p<0.001, ****p<0.0001; ns, not significant.
